# CD39^+^ conventional CD4^+^ T cells with exhaustion traits and cytotoxic potential infiltrate tumors and expand upon CTLA-4 blockade

**DOI:** 10.1101/2023.01.30.526262

**Authors:** Sabrina N. Bossio, Carolina Abrate, Jimena Tosello Boari, Constanza Rodriguez, Fernando P. Canale, María C. Ramello, Wilfrid Richer, Dario Rocha, Christine Sedlik, Anne Vincent-Salomon, Edith Borcoman, Andres Del Castillo, Adriana Gruppi, Elmer Fernandez, Eva V. Acosta Rodríguez, Eliane Piaggio, Carolina L. Montes

**Affiliations:** Departamento de Bioquímica Clínica, Facultad de Ciencias Químicas, Universidad Nacional de Córdoba, Córdoba, Argentina; Centro de Investigaciones en Bioquímica Clínica e Inmunología (CIBICI-CONICET), Córdoba, Argentina; PSL Research University, Institut Curie Research Center, Translational Research Department, Paris, France; INSERM U932, Paris, France; Centro de Investigación y desarrollo en inmunología y enfermedades infecciosas (CIDIE-CONICET); Diagnostic and Theranostic Medicine Division, Institut Curie, PSL Research University, Paris, France; Department of Medical Oncology, Institut Curie, Paris, France; Hospital Rawson, Polo Sanitario, Córdoba, Argentina

**Keywords:** CD39, Conventional CD4^+^ T cells, exhaustion, cancer, cytotoxicity

## Abstract

**Background:** Conventional CD4^+^ T (Tconv) lymphocytes play important roles in tumor immunity; however, their contribution to tumor elimination remains poorly understood.

**Methods:** Here we describe a subset of Tconv cells characterized by the expression of CD39. The phenotype, the effector function and transcriptional profile of tumor-infiltrating CD39^+^ Tconv lymphocytes from different mouse cancer models and breast cancer patients were studied by multiparametric flow cytometry and RNA sequencing. The impact of the *in vivo* CTLA-4 blockade on the tumor-infiltrating CD39^+^ Tconv population was assessed in mice grafted with the immunogenic MC38 colorectal tumor. Overall survival was evaluated in a cohort of patients from the TCGA consortium.

**Results:** In mouse cancer models, we observed that CD39^+^ Tconv cells accumulated in tumors as they grew but were absent in lymphoid organs. Compared to tumor CD39^−^ counterparts, CD39^+^ Tconv cells exhibited a cytotoxic and exhausted signature at the transcriptomic level, confirmed by high protein expression of inhibitory receptors and transcription factors related to the exhaustion phenotype. Additionally, CD39^+^ Tconv cells showed increased production of IFN gamma, granzyme B, perforin and CD107a expression, but reduced production of TNF. *In vivo* CTLA-4 blockade induced the expansion of tumor CD39^+^ Tconv cells, which maintained their cytotoxic and exhausted features. In breast cancer patients, CD39^+^ Tconv cells were found in tumors and in metastatic lymph nodes but were less frequent in adjacent non-tumoral mammary tissue and not detected in non-metastatic lymph nodes and blood. Human tumor CD39^+^ Tconv cells constituted a heterogeneous cell population with features of exhaustion, impaired TNF production, and high expression of inhibitory receptors and CD107a. We found that high CD4 and ENTPD1 (CD39) gene expression in human tumor tissues correlated with a higher overall survival rate in breast cancer patients.

**Conclusions:** We found that CD39 acts as a biomarker of Tconv cells with characteristics of both exhaustion and cytotoxic potential. CTLA-4 blockade expands CD39^+^CD4^+^ T cells which may contribute to the reduction of tumor development. Discovering the role of CD39-expressing CD4^+^ T cells in the tumor microenvironment should help design new strategies to manipulate them and improve the efficacy of current immunotherapies.

## Introduction

CD8^+^ T cells are fundamental players in the anti-tumor immune response. The role of CD4^+^ Tconv cells in controlling or enhancing tumor response is less understood. However, antigen-specific Tconv cells accumulate at tumor sites, supporting their involvement in anti-tumor effector functions, in both mouse experimental models and human cancer ^1, 2^.

Antigen-driven activation of naïve CD4^+^ T cells leads to their differentiation into effector T cell subsets ^3^. Differentiated CD4^+^ T cells (e.g. Th1, Th2, Th17, and regulatory T cells) impact tumor immunity by influencing other cell types ^4^. A less described cytotoxic activity has been observed in a subset of CD4^+^ T cells (cytotoxic CD4^+^ T cells, CTLs) ^5^. CD4^+^ CTLs were identified in murine cancer models, where melanoma-specific CD4^+^ T cells gained cytotoxic activity and eradicated tumors ^6^. Similarly, NY-ESO1-specific CD4^+^ T cells from melanoma patients can lyse melanoma cells in a MHCII-dependent fashion ^7^.

Within the tumor microenvironment (TME), CD4^+^ T cell populations have been described with effector, regulatory, and exhaustion features ^8, 9^. T cell exhaustion was initially described in CD8^+^ T lymphocytes, in animal models and in patients with cancers or chronic viral infections ^10-12^. Two features of CD8^+^ T cell exhaustion are the loss of effector cytokine production, and the co-expression of inhibitory receptors (iRs) ^13^. Exhausted CD8^+^ T cells display a particular pattern of transcription factors (TFs), including T-bet, Eomes, Blimp-1, and TOX ^14, 15^. Recently we described that CD39, an ecto-nucleotidase involved in adenosine production, defines cell exhaustion in human and mouse tumor-infiltrating (T-I) CD8^+^ T cells ^16^. In addition, it has been demonstrated that CD39 identifies tumor-reactive CD8^+^ T cells in human solid tumors ^17^.

CD39 is constitutively expressed in Treg cells ^18^; it has been demonstrated that T-I PD-1^high^CD39^+^ Tconv cells from head and neck, ovarian, and cervical cancer patients exhibit features of T cell exhaustion ^19^. CD39 also identifies a tumor-specific CD4^+^Foxp3^−^ T cell population in squamous cell carcinoma patients ^20^.

Here, we describe that CD39^+^ Tconv infiltrate tumors from mouse cancer models and breast cancer (BC) patients. Using phenotypic and transcriptional approaches, we demonstrate that CD39^+^ Tconv exhibit features of exhaustion and transcriptional signature of cytotoxic T cells. Moreover, high expression of CD4 and ENTDP1 (CD39) in tumors correlates with better survival in BC patients. Together, our findings reinforce the idea that the CD39^+^ Tconv population likely plays a significant role in the immune response against tumors.

## Materials and Methods

### Mouse and human samples

Mice (6–10 weeks) were housed at the animal facility of the CIBICI-CONICET. Animal protocols were approved by the Institutional Animal Care and Use Committee (IACUC) of the CIBICI-CONICET.

Human tumors, juxtatumoral breast tissues (adjacent non-tumoral mammary tissue), and tumor-draining lymph nodes (dLNs) were collected from 49 untreated female BC patients undergoing standard-of-care surgery at the Institut Curie Hospital (France) and the Hospital Rawson (Argentina). BC patients had no treatment prior to surgery and without chronic infections. Clinical and pathological data of recruited/analyzed BC patients are described in Supplementary Table 1. All protocols were approved by the institutional review boards and all patients signed an informed consent form. dLNs were classified by histology into metastatic or non-metastatic according to the presence of tumor cells, and confirmed by Epcam/CD45 staining using flow cytometry. Blood was sampled from patients of Hospital Rawson. All studies in patients were conducted in accordance with the ethical guidelines of the Declaration of Helsinki.

Tumors and dLNs were disaggregated mechanically and enzymatically with liberase and DNase I (Roche). Peripheral blood mononuclear cells (PBMCs) were isolated by centrifugation over Ficoll-Paque gradients (GE Healthcare).

### Cell lines

MC38 and MCA-OVA were kindly provided by Dr. Clothilde Théry. 4T1 and B16F10-OVA cell lines were purchased from ATCC. B16F10-OVA and MC38 were maintained in DMEM (GIBCO). 4T1 and MCA-OVA in RPMI-1640. Media were supplemented with 10% Fetal Bovine Serum (GIBCO), 1mM L-Glutamine (GIBCO), 25 mmol/L Hepes (Cell Gro), and 40 ug/mL gentamicin sulfate (Richet).

### *In vivo* tumor models

Male C57BL/6 or Foxp3-EGFP reporter mice were inoculated subcutaneously (s.c.) with 1×10^6^ B16F10-OVA, 0.5×10^6^ MC38, or 0.5×10^6^ MCA-OVA cells. Female Balb/c mice were inoculated s.c. with 3×10^4^ 4T1 cells. Tumors excised at day indicated in each figure were disaggregated mechanically and enzymatically with 2 mg/mL collagenase IV and 50 U/mL DNase I (Roche).

### Anti-CTLA-4 blockade

Male C57BL/6 mice were inoculated s.c. with 0.5×10^6^ MC38 cells. Tumor-bearing mice were treated with anti-CTLA-4 (BioXCell) (n=8) or control IgG (n=8) (Syrian hamster IgG BioXCell), following the protocol depicted in Fig 4A.

### Flow Cytometry

Single cell suspensions were stained with monoclonal antibodies against mouse and human antigens. For fluorochromes and clones, refer to Supplementary Table 2. Gating strategies for cellular populations from mouse and human tissues are shown in Supplementary Figure 1.

TFs (intracellular staining) and intracellular cytokine detection cells were performed following protocols previously described ^16, 21^.

Samples were acquired in a BD LSR Fortessa flow cytometer (BD Biosciences) and the data analyzed with FlowJO software.

### Cell sorting and RNA isolation for RNA sequencing

B16F10-OVA tumors from 9 Foxp3-EGFP tumor-bearing mice were processed for RNA sequencing. Tumors were shaped into 3 pools, and purified CD4^+^ T cells were obtained by positive selection with the “EasySep Mouse CD4 Positive Selection II” kit (StemCell Technologies). Then, cells were stained with CD45-AF700, TCRβ-APC-Cy7, CD8-APC, CD4-PE-TexasRed, CD39-PE, CD11b-PE-Cy7, CD19-PE-Cy7, NK1.1-PE-Cy7, F4/80-PE-Cy7, and later with 4’,6-diamidino-2-phenylindole (DAPI) LIVE/DEAD stain. Among the live CD4^+^ T cells (DAPI- CD11b- CD19- NK1.1- F4/80- CD8- CD45+ TCRβ+ CD4+), an equal number of Treg, CD39 positive Tconv, and CD39 negative Tconv were sorted with a purity of 98-99%, using a BD FACS ARIA II cell sorter (GFP+, GFP- CD39+, GFP- CD39-, respectively). Cells were lysed with TCL buffer (Qiagen) with 1% of ß-mercaptoethanol and stored at -80°C.

RNA was isolated using a Single Cell RNA purification kit (Norgen), including RNase-Free DNase Set (Qiagen) treatment. RNA integrity numbers were evaluated with an Agilent RNA 6000 pico kit. Samples were assessed following the manufacturer’s instructions.

### RNA sequencing

Reverse transcription and cDNA amplification were performed with the SMART-Seq v4 Ultra Low Input RNA Kit (Takara). Barcoded Illumina-compatible libraries were generated from 5 to 10 ng of DNA of each sample, using the Nextera XTP Preparation Kit (illumina). Libraries were sequenced on an Illumina Novaseq 6000 using 100Lbp paired-end mode, 20 million reads per sample.

### RNA analysis

FASTQ files were mapped to the ENSEMBL Mouse (GRCm38/mm10) reference using STAR 2.3 ^22^ (STAR, RRID:SCR_004463) and counted by the feature Counts from the RSubread R package. Read count normalization and group comparisons were performed with EdgeR (edgeR, RRID:SCR_012802) and limma R libraries. Genes in which total sum counts per million were >10 were kept for differential expression analysis. Differential expression between experimental conditions was defined by using the “treat” limma method, using p-value ≤ 0.05 and fold change ≥ 2. Volcano plots were made with R Version +3 and imaged by Java Treeview software. Functional analyses were performed with GSEA and the Panther Classification System. GEO Accession ID : GSE21881.

### TCGA data

The Cancer Genome Atlas (TCGA) ^23^ data was downloaded with TCGA assembler ^24^. Raw counts and transcripts per million (TPM) were obtained for expression data. Normalization factors for raw count data were calculated by Trimmed Mean of M-values (TMM) with the EdgeR package ^25^. Raw count expression data was transformed to log_2_ (counts per million+0.5).

### Survival analysis

The expression of PTPRC, CD4, and FOXP3 genes was categorized as “high” or “low” according to the median. Samples with high PTPRC, high CD4, and low FOXP3 expression were selected for analysis. CD39 expression was categorized into “high” or “low” according to a cutpoint defined by maximizing the log-rank statistic of a Kaplan-Meier model for overall survival, using the surv_cutpoint function of the survminer R package ^26^.

### Statistical analysis

GraphPad Prism 8.0 software was used for statistical analysis. In experiments with mouse and human samples, a paired Student’s *t*-test was used to compare groups. One-way ANOVA with the Tukey post-test was used for multiple comparisons. *P* values <0.05 were considered statistically significant. Z-score was estimated as: (MFI-MFI_average_)/SD.

## Results

### CD39^+^ Tconv cells infiltrate tumors from mouse experimental models

We first explored the presence of CD4^+^FOXP3^−^ conventional T cells expressing CD39 in tumor, spleen, and tumor-draining lymph nodes (dLNs) from B16F10-OVA tumor-bearing mice. On day 17 post injection (p.i.), we observed that around 50% of T-I Tconv expressed CD39. In contrast to tumor, CD39^+^ Tconv were nearly absent in dLNs and spleen (Fig. 1A). Low frequencies of this cell subset were detected in dLNs (1.63±0.23) and spleen (4.18±1.08) from tumor-free mice. As observed in the B16F10-OVA model, Tconv from mice bearing MCA-OVA, MC38 or 4T1 tumors also showed a high frequency of CD39^+^ cells (Supplementary Fig. S2 A-C), while, in lymphoid organs, CD39^+^ Tconv cells were present at significantly lower frequencies than in tumors. This indicates that CD39 expression on Tconv cells is associated with the TME, and that CD39 expression in T-I Tconv is a common feature across tumors of different histological origin.

**Figure 1.**
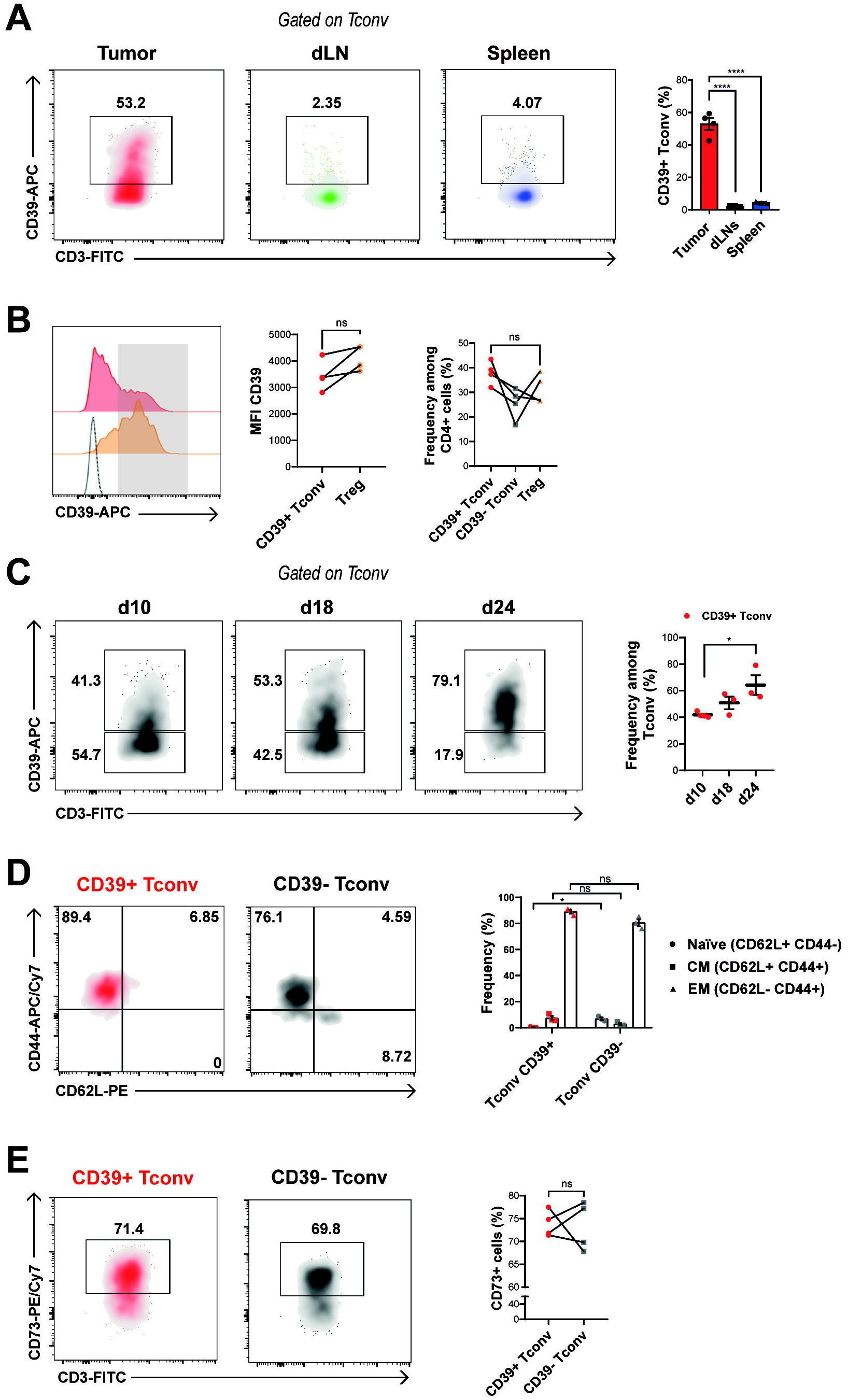
CD39^+^ Tconv infiltrate tumors from B16F10-OVA tumor-bearing mice. **A** Representative density plots and graphs show frequency of CD39^+^ cells gated on Tconv cells (CD4^+^FOXP3^−^) from tumors, dLN and spleen at day 17 p.i. (for gating strategy see Supplementary Figure S1 A and B). **B** (Left and middle panels) Histograms and line graphs show CD39 expression on Treg (orange histogram and dots) or Tconv cells (red histogram and dots). (Right panel) Frequency of T-I CD39^+^ Tconv (red dots), CD39^−^ Tconv (gray squares) and Treg (orange triangles) among CD4^+^ cells. **C** Frequency of CD39^+^ cells gated on T-I Tconv at different timepoints during tumor progression. Mean values and statistical analyses (day 10 vs. day 24) are shown. **D** Representative density plots and graphs show the frequency of cells expressing CD44 and CD62L within CD39^+^ or CD39^−^ T-I Tconv cells at day 17 p.i. **E** Representative density plots and line graphs show frequency of CD73-expressing CD39^+^ or CD39^−^ T-I Tconv cells at day 17 p.i. All results are representative of 3 to 5 independent experiments. **A-E:** data presented as mean ± SEM. ns, non-significant; * *p*≤ 0.05; \*\**p*≤ 0.01; or **** *p*≤ 0.001. **B, E** Lines indicate that data are paired. Paired T test was used to compare CD39^+^ vs CD39^−^ Tconv cells.

Considering that CD39 is highly expressed by Treg as part of their immunosuppressive arsenal, we compared CD39 expression in T-I Tconv and Treg from B16F10-OVA tumor-bearing mice. We observed that CD39 expression, determined as MFI, in Treg was similar to that observed in CD39^+^ Tconv (Fig. 1B, left panel). Also, we found similar frequencies of CD39-expressing Tconv and Treg (Fig.1B, right panel).

A kinetic study in B16F10-OVA tumors showed that the frequency of CD39^+^ Tconv increased with tumor progression. Indeed, the percentage of T-I CD39^+^ Tconv was significantly higher at day 24 p.i than at day 10 p.i. (Fig. 1C). The expression of CD39 in T-I Tconv thus correlates with tumor progression.

We further characterized the activation/differentiation phenotype of T-I CD39^+^ Tconv and found that most of these cells exhibited an effector memory (EM) phenotype (CD62L^−^CD44^+^). We found naïve cells (CD62L^+^CD44^−^) only among CD39^−^ Tconv (Fig.1D). These results suggest that, within the TME, Tconv cells, including CD39^+^ T cells, have probably undergone antigenic stimulation.

Like Treg ^18^, around 70% of Tconv also expressed CD73, regardless of CD39 expression. CD73 is an enzyme that works in cooperation with CD39 in converting extracellular ATP to adenosine (Fig. 1E). These findings suggest that Tconv could have a role in purinergic signaling in the TME, with CD39^+^Tconv being an effector population that can generate adenosine by itself.

### CD39^+^ Tconv shows both a cytotoxic and exhaustion transcriptomic signature

To compare the transcriptional landscape between different CD39-expressing CD4^+^ T-cells (Tconv and Treg) and to understand the phenotypic differences between CD39^+^ vs CD39^−^ Tconv, we isolated three replicates of T-I CD4^+^ T cell subsets.

Using Foxp3-EGFP tumor-bearing mice we obtained CD4^+^GFP^+^ (Treg), CD4^+^CD39^−^ GFP^−^ (CD39^−^ Tconv), and CD4^+^CD39^+^GFP^−^ (CD39^+^ Tconv) from tumors by cell sorting (Fig. 2A). Principal component analysis (PCA) showed that the three CD4^+^ T cell subsets clustered separately, reflecting their different transcriptional profiles (Fig. 2B). To define the molecular profile of CD39^+^ Tconv, we searched for the genes differentially expressed (DEGs) between CD39^+^ Tconv and CD39^−^ Tconv cells and determined that 449 and 153 genes were significantly up- or downregulated, respectively (Fig. 2C). Those upregulated in CD39^+^ Tconv included genes associated with cytotoxicity (such as *Gzmb, Gzmc, Gzmf, Prf1, Nkg7, Eomes)*, and with T cell exhaustion (such as *Lag3, Havcr2 (*Tim-3*), Tigit, Cd160, Pdcd1 (*PD-1) and *Tox*) (Fig. 2C). CD39^+^ Tconv also upregulated genes corresponding to chemokines (such as *Cxcl1, Ccl3, Ccl4 and Ccl5*), or genes characteristic of tissue-resident T cells (such as *Itga1 (*CD49a*), Cxcr6 and Alox5ap*), among others (see Supplementary Table 3). CD39^+^ Tconv displayed a reduced expression of *Tcf7* (Tcf-1) (Fig. 2C), a common feature of terminally differentiated exhausted T cells that contrasts with Tcf-1 expression in cells with stem cell-like properties akin to memory T cell populations ^27^.

**Figure 2:**
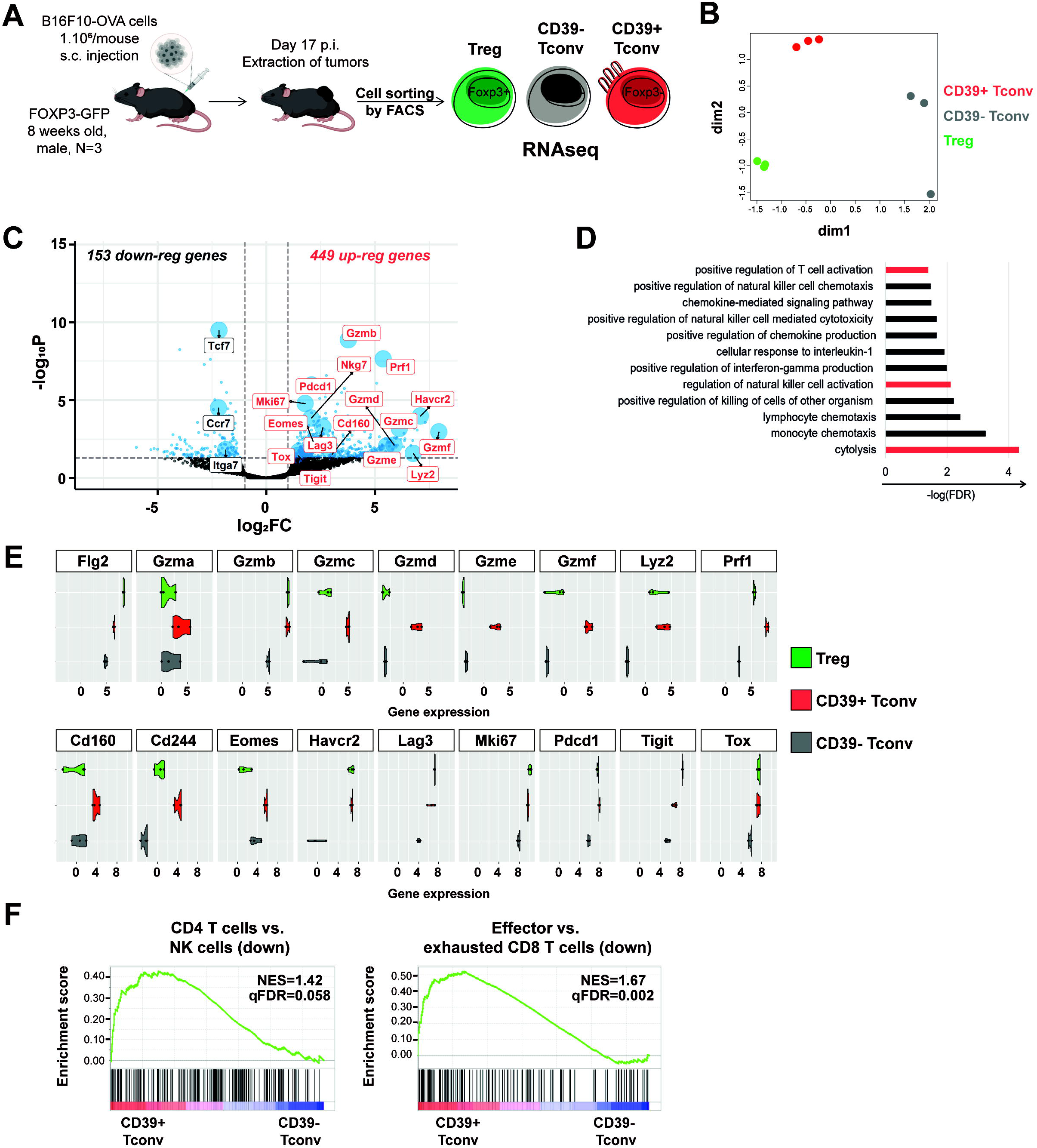
Transcriptional profiling of tumor-infiltrating Tconv cells. **A** Scheme of sort strategy for B16F10-OVA T-I Tconv and Treg cell isolation for transcriptional analysis from FOXP3-EGFP mice (N=3). **B** Multidimensional scaling (MDS) two-dimensional (dim1 and dim2) scatterplot, where distances on the plot approximate the typical log2 fold changes between all the analyzed samples. Each dot represents a sample, colored by subset. **C** Volcano plot showing differentially expressed genes (DEGs) between CD39^+^ Tconv and CD39^−^ Tconv cells. Genes were considered differentially expressed if FC≥ 2 and adjusted p value<0.05, and these are colored in blue. Most relevant up- and downregulated genes are shown in red and gray, respectively, and are highlighted with their gene symbols. **D** Graph shows selected pathways significantly enriched (p<0.05) corresponding to genes upregulated in CD39^+^ Tconv compared to CD39^−^ Tconv cells using the Panther Classification System. Relevant pathways are colored in red. **E** Violin plots show the gene expression values (voom normalized log-cpm values in log2 scale) related to cytotoxicity and T cell exhaustion on Treg (green), CD39^+^ Tconv (red), and CD39^−^ Tconv (gray). **F** GSEA plots show selected gene sets (GSE27786-left- and GSE9650-right-) enriched in CD39^+^ Tconv vs CD39^−^ Tconv cells. Normalized enrichment score (NES) and q False Discovery Rate (qFDR) are indicated in the graph. \**p*≤ 0.05. P values were calculated using paired T test.

Biological pathways enriched in CD39^+^ Tconv included: “positive regulation of T cell activation”, “lymphocyte chemotaxis”, “positive regulation of interferonγ production”, “chemokine-mediated signaling pathway”, as well as pathways and processes associated with NK activation, such as “cytolysis”, “regulation of NK activation”, “positive regulation of NK cell mediated cytotoxicity”, and “positive regulation of NK cell chemotaxis” (Fig. 2D). Compared to CD39^−^ Tconv, CD39^+^ Tconv showed higher gene expression of genes related to cytotoxicity (*Flg2, Gzms, Lyz2* and *Prf1*) and T-cell exhaustion (such as iRs, *Tox, Eomes* and *Mki67*) (Fig 2E). Gene set enrichment analysis (GSEA) indicated that the transcriptional signature of CD39^+^ Tconv was enriched in genes associated with NK-signature and CD8^+^ T cell exhaustion compared to CD39^−^ Tconv cells (Fig. 2F). Overall, the transcriptional analysis revealed that T-I CD39^+^ Tconv from B16F10-OVA tumor-bearing mice are enriched in genes associated with T cell activation, cytotoxicity, and exhaustion compared to CD39^−^ Tconv cells.

Since the expression of CD39 and CD73 suggests a regulatory role in CD39^+^ Tconv, we explored the core differences between Foxp3^+^ and Foxp3^−^ CD39^+^ CD4^+^ T cells, comparing T-I CD39^+^ Tconv and Treg transcriptomes. Differential expression analysis highlighted that 244 genes were significantly upregulated and 203 genes significantly downregulated in CD39^+^ Tconv compared to Treg. As expected, CD39^+^ Tconv exhibited lower levels of genes typically associated with Tregs, such as *Foxp3, Il10, Ctla4, Il2ra*, and *Ccr8*, among others (see Supplementary Fig. S3A and Supplementary Table 4). Notably, compared to Treg, CD39^+^ Tconv upregulated cytotoxicity-related genes, such as *Gzmd, Gzme, Gzmf, Crtam, Serpinb9*, and *Prf1* (Supplementary Fig. S3A). Additionally, CD39^+^ Tconv showed enrichment in pathways associated with lymphocyte activation, cell killing, cytolysis, and regulation of natural killer cell activation, among others (Fig. Supplementary S3B). These results indicate that CD39^+^ Tconv may have effector functions.

### The phenotype of tumor-infiltrating CD39^+^ Tconv is associated with exhaustion and cytotoxic potential

We next evaluated the protein expression of iRs and TFs associated with T cell activation and exhaustion in T-I CD4^+^ T cells from B16F10-OVA tumor-bearing mice. Compared with their CD39^−^ Tconv counterparts, T-I CD39^+^ Tconv population exhibited the highest frequency of cells expressing all the evaluated iRs (PD-1, TIGIT, Tim-3, LAG-3, and 2B4) (Fig. 3 A, Left panel). CD39^+^ Tconv exhibited higher mean fluorescence intensity (MFI) of PD-1, TIGIT, Tim-3, LAG-3, 2B4, CTLA-4, and PDL-1 than CD39^−^ Tconv (Supplementary Fig. S4A and B). They also showed a higher proportion of iR co-expression, with a higher frequency of T cells expressing 4 or 5 iRs (Fig. 3 A, Right panel). Moreover, T-I CD39^+^ Tconv exhibited higher expression, determined as MFI, of TFs, such as T-bet, Helios, Eomes, Blimp-1, cMaf, Tox, and Ki67, but lower expression of TCF-1 than CD39^−^ Tconv (Fig 3B). Overall, matching the transcriptional data, the protein expression of CD39 is paralleled by high expression of iRs and TFs associated with effector function and exhaustion, as well as lower TCF-1 expression.

**Figure 3:**
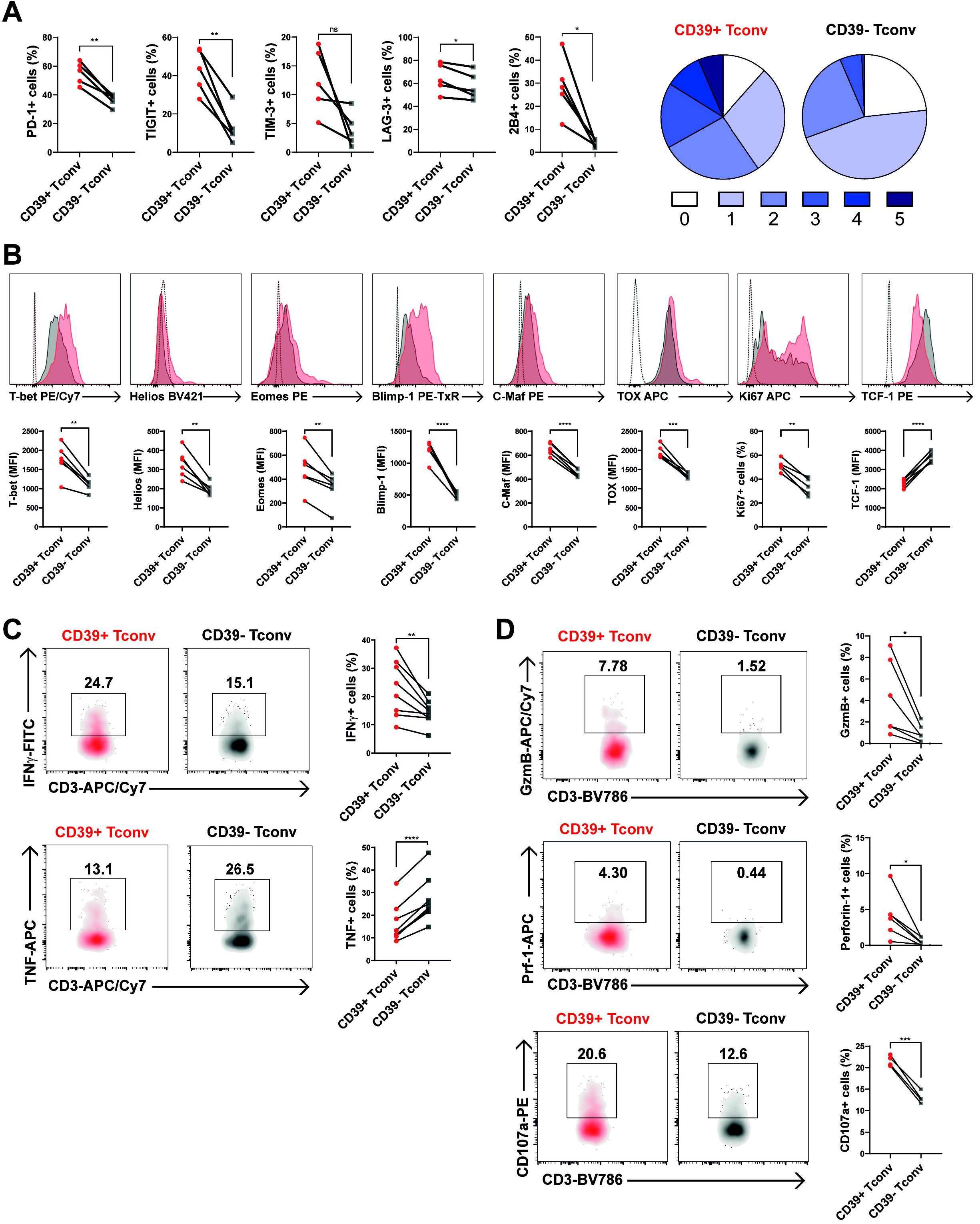
T-I CD39^+^ Tconv cells exhibit a phenotype of exhaustion and cytotoxicity. T-I lymphocytes were obtained from tumors from B16F10-OVA tumor-bearing mice at day 17 p.i. and analyzed by flow cytometry. **A** (Left panel) Line graphs show frequencies of iR-expressing CD39^+^ (red dots) or CD39^−^ (gray squares) T-I Tconv cells. (Right panel) Pie charts show the mean proportion of cells expressing zero to five of the evaluated iRs (PD-1, TIGIT, TIM-3, LAG-3 and 2B4) in CD39^+^ or CD39^−^ T-I Tconv cells. **B** Representative histograms and line graphs show the expression of selected TFs on CD39^+^ Tconv (red histograms, red dots) and CD39^−^ Tconv (gray histograms, gray squares). Dotted lines represent unstained controls. **C** Representative density plots and line graphs show the frequency of IFNγ^+^ or TNF^+^ cells among CD39^+^ (red density plots, red dots) or CD39^−^ (gray density plots, gray squares) T-I Tconv cells after PMA/Ionomycin stimulation. **D** Representative density plots and line graphs show the frequency of granzyme B^+^, perforin^+^, or CD107a^+^ cells among CD39^+^ (red density plots, red dots) or CD39^−^ (gray density plots, gray squares) T-I Tconv cells after PMA/Ionomycin stimulation. All results are representative of 3 to 5 independent experiments. **A-D** Lines indicate that data are paired. Paired T test was used to compare CD39^+^ vs CD39^−^ Tconv cells. \**p*≤ 0.05; \*\**p*≤ 0.01; \*\*\**p*≤ 0.001; \*\*\*\**p*≤ 0.0001

To further evaluate CD39^+^ Tconv effector functions, we evaluated by FACS the frequency of cytokine production within the Tconv population. CD39^+^ as well as CD39^−^ Tconv were able to produce IFNγ. Surprisingly, although CD39^+^ Tconv exhibited a higher percentage of IFNγ^+^ cells, their TNF production was lower than CD39^−^ Tconv (Fig. 3C). Furthermore, in accordance with their IFNγ production phenotype and T-bet expression, CD39^+^ Tconv exhibited higher cytotoxic potential than CD39^−^ counterpart, as attested by intracellular granzyme B and perforin expression together with the CD107a mobilization after *ex vivo* stimulation (Fig. 3D).

Overall, T-I CD39^+^ Tconv represent a distinct CD4^+^ T cell population, with transcriptomic signatures and phenotypic markers of activation/proliferation and cytotoxicity, but also exhaustion.

### Anti-CTLA-4 blockade induces the expansion of TI-CD39^+^ Tconv

Anti-checkpoint blockade acts on distinct T lymphocyte subsets, with anti-PD-1 predominantly inducing the expansion of T-I exhausted CD8^+^ T cells, while anti-CTLA-4 treatment preferentially expands Th1-like CD4^+^ effector cells and reactivates exhausted CD8^+^ T cells ^28^. We therefore investigated whether CTLA-4 blockade affects the TI-CD39^+^ Tconv population in mice grafted with the immunogenic MC38 colorectal tumor. Tumor-bearing mice were treated with anti-CTLA-4 or control IgG, as indicated in Fig 4A, following a previously described treatment schedule ^28^. As expected, tumor growth was reduced in MC38 tumor-bearing mice with anti-CTLA-4 treatment (Supplementary Figure S5). In agreement with previous reports ^29^, anti-CTLA-4 treated mice showed higher frequencies of tumor CD8^+^ T cells, reduced frequencies of Treg, and no change in Tconv frequencies among total CD3^+^ T cells, compared to IgG control-treated mice (Fig.4B). Focusing on T-I CD4^+^ T cell subpopulations, we found that anti-CTLA-4 induced an increase in CD39^+^ Tconv frequencies (Fig.4C). Notably, the expression by these expanded CD39^+^ Tconv cells of all the iRs evaluated except LAG-3 (Fig. 4D), as well as of TFs related to exhaustion (Eomes, TOX, Helios), was unchanged with respect to CD39^+^ Tconv from IgG-treated mice. Furthermore, the T-I CD39^+^ Tconv subset from anti-checkpoint treated mice showed increased frequencies of cells expressing T-bet and conserved frequencies of cells able to mobilize CD107a, compared to control mice (Fig.4E).

**Figure 4:**
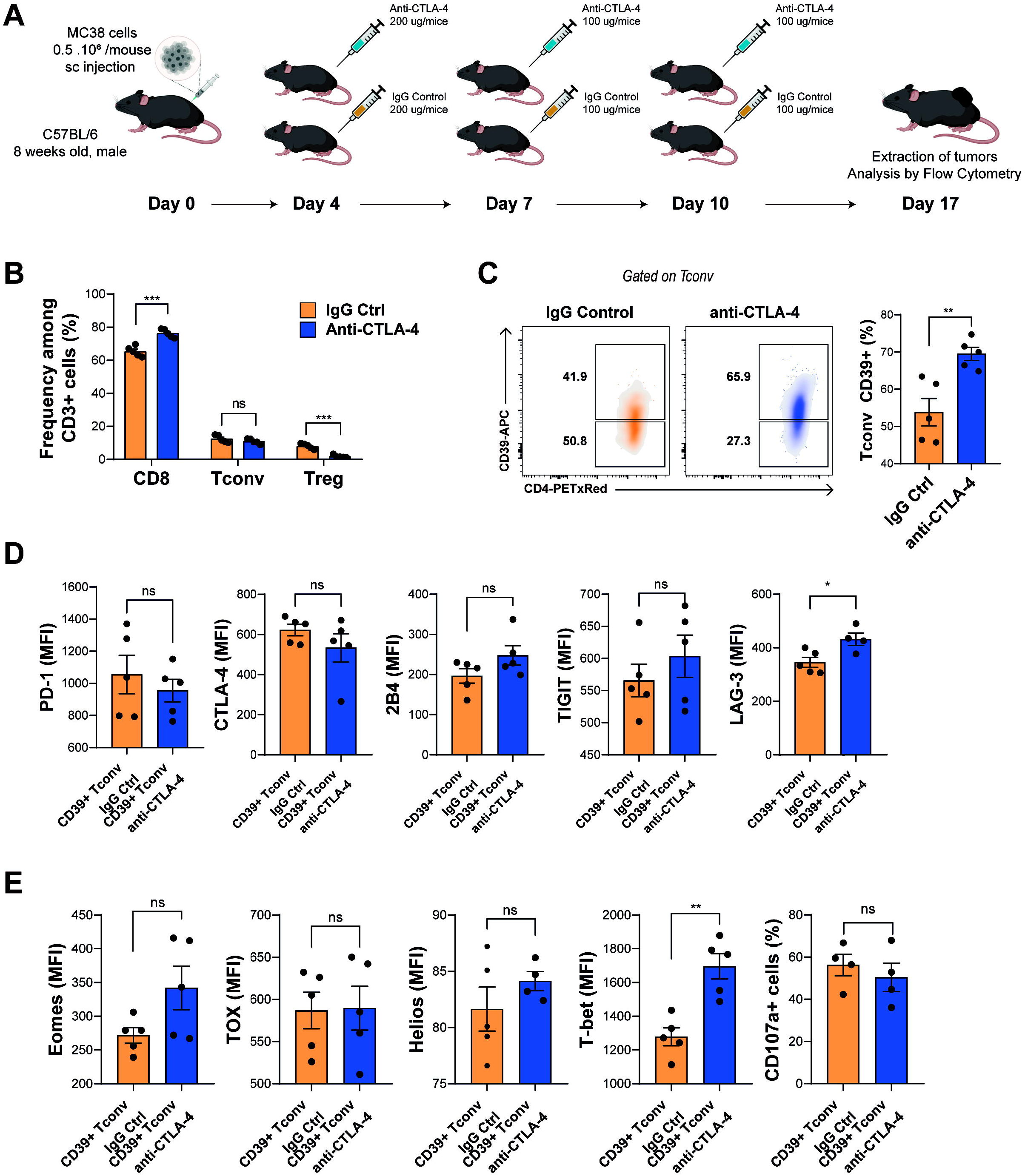
Anti-CTLA-4 treatment induces the expansion of T-I CD39^+^ Tconv cells. **A** Experimental design of anti-CTLA-4 treatment of MC38 tumor-bearing mice. **B** Graphs show frequency of T-I CD8^+^, Tconv and Treg cells in mice treated with anti-CTLA-4 (blue) or treated with IgG control (orange). **C** Representative density plots and graphs show the frequency of CD39^+^ Tconv cells from tumors of MC38 tumor-bearing mice treated with anti-CTLA-4 (blue) or IgG control (orange). **D** Bar graphs show the expression as MFI of iRs (PD-1, CTLA-4, 2B4, TIGIT, LAG-3) on T-I CD39^+^ Tconv cells from mice treated with anti-CTLA-4 (blue) or IgG control (orange). **E** Bar graphs show the expression as MFI of selected TFs (Eomes, TOX, Helios and T-bet) on T-I CD39^+^ Tconv from tumors of treated with anti-CTLA-4 (blue) or IgG control (orange) and frequency of CD107a+ T-I CD39^+^ Tconv cells from anti-CTLA-4 (blue) or IgG control (orange) treated mice. All results are representative of 2 independent experiments. **B-E** Data presented as mean ± SEM. P values were calculated using unpaired T test. ns, non-significant, *p≤ 0.05 **p≤ 0.01.

These results indicate that anti-CTLA-4 treatment induces the expansion of T-I CD39 expressing Tconv which, despite maintaining exhaustion features, conserve their cytotoxic potential and may contribute to the reduction of tumor development.

### CD39^+^ Tconv cells accumulate within tumors and metastatic lymph nodes from breast cancer patients

Understanding the role of T-I CD4^+^ T cells in cancer patients may be key for prognosis evaluation and for the improvement of therapy design ^30^. We therefore evaluated the distribution of the CD4^+^ T cell population (Treg and Tconv) in samples from 12 primary BC tumors matched with juxtatumoral breast tissue (adjacent non-tumoral mammary tissue). As previously described ^31^, BC tumors had a higher Treg infiltration than juxtatumoral tissues, and here we observed that the relative percentage of Tconv was significantly lower in the tumor niche than in juxtatumoral tissue (Fig. 5A left panel). Nevertheless, Tconv was still the major CD4^+^ subset in both tissues. Among total Tconv, CD39^+^ cells were present at a sizable frequency both in tumoral and juxtatumoral tissues (6.07±3.70% and 4.32±4.12%, respectively), although the percentage was higher in tumors (Fig.5A right panel). No CD73 expression was detected in T-I CD39^+^ Tconv (data not shown). These results suggest that the TME may promote the accumulation of Treg and Tconv expressing CD39.

**Figure 5.**
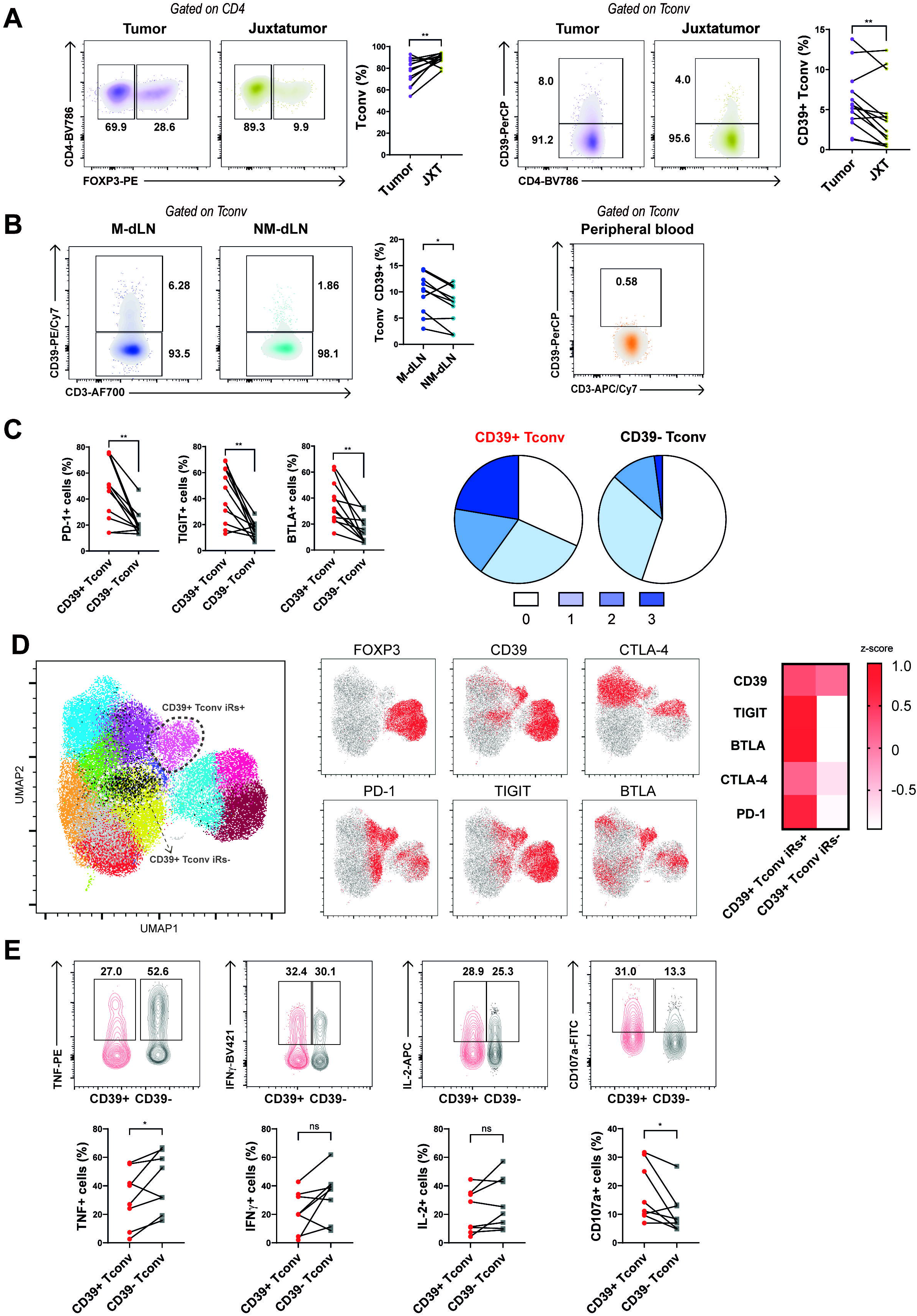
CD39^+^ Tconv cells infiltrate human breast tumors and metastatic lymph nodes and exhibit high expression of iRs. **A** (Left panel) Representative dot plots and line graph show frequencies of Tconv cells from tumors (violet density plot and dots) and juxtatumoral tissues (yellow density plot and dots) from BC patients (N=13). (Right panel) Representative dot plots and line graph show frequencies of CD39^+^ cells gated on Tconv from tumors (violet density plot and dots) and juxtatumoral tissues (yellow density plot and dots) from BC patients (N=13). **B** (Left panel) Representative dot plots and line graph show frequencies of CD39^+^ Tconv cells from matched M-dLNs (blue density plot and dots) and NM-dLNs (dark blue density plot and dots) from BC patients (N*=9)*. (Right panel) Representative dot plot shows frequency of CD39^+^ Tconv cells in PB from BC patients (one out of seven samples) (gating strategies are showed in Supplementary Figure S1). **C** (Left panel) Line graphs show frequencies of iR-expressing CD39^+^ (red dots) or CD39^−^ (gray squares) T-I Tconv cells from BC patients. (Right panel) Pie charts show the mean proportion of cells expressing zero to 3 of the evaluated iRs (PD-1, TIGIT and BTLA) in CD39^+^ or CD39^−^ T-I Tconv cells. **D** (Left panel) UMAP projection of 51,100 T-I CD4^+^ cells from BC patients (N=7). Colors represent different clusters within CD4^+^ cells defined by common phenotypic markers (CD45RA, CD39, CD27, FOXP3, TIM-3, TIGIT, BTLA, CTLA-4, PD-1, LAG-3). (Middle panel) UMAP projections show cells colored according to the expression of different markers: positive for FOXP3, CD39, CTLA-4, PD-1, TIGIT, and BTLA in red, and negative for those in gray. (Right panel) Heat map shows normalized expression of iRs related to exhaustion from 2 clusters highlighted as *Tconv CD39+iRs+* and *Tconv CD39+iRslow/-*. **E** Representative dot plots and line graphs show frequencies of cytokine-producing cells (IL-2, TNF and IFNγ) and CD107a^+^ cells in CD39^+^ (red density plots and dots) and CD39^−^ (gray density plots and squares) T-I Tconv cells from BC patients after PMA/Ionomycin stimulation (*N=*7). **A, B, C, E** Lines indicate that data are paired. P values were calculated using paired T test. *p≤ 0.05, **p≤ 0.01, ns, non-significant.

Considering that tumor-specific cells may be primed in the dLNs and more importantly in metastatic dLNs (M-dLNs) where tumor antigens are highly abundant, we evaluated the presence of CD39^+^ Tconv cells in metastatic and non-metastatic dLNs and peripheral blood (PB) from BC patients. In agreement with our observations regarding T-I CD8^+^ T cells ^16^, M-dLNs had higher frequencies of CD39^+^ Tconv cells among total CD3^+^ cells than non-metastatic draining lymph nodes (NM-dLNs). In addition, CD39^+^ Tconv cells were not detectable in PB (Fig 5B).

We further evaluated the differential distribution of iRs in patient-matched CD39^+^ or CD39^−^ T-I Tconv cells. CD39^+^ Tconv showed significantly higher frequencies of PD-1-, TIGIT- and BTLA-expressing cells (Fig. 5C Left panel), as well as more frequency of cells co-expressing two or three iRs (Fig. 5C Right panel). Similarly, CD39^+^ Tconv cells from M-dLNs showed higher frequency of PD-1, TIGIT and BTLA-expressing cells than CD39^−^ Tconv (Supplementary Fig. S6A).

To define the phenotype of CD39^+^ Tconv in the tumor, we applied high-dimensional flow cytometry analysis to our data. Unsupervised analysis using the dimension reduction algorithm, Uniform Manifold Approximation and Projection (UMAP), followed by the Phenograph clustering algorithm, identified 14 clusters (Figure 5D). Three clusters corresponded to Treg cells, as indicated by the expression of Foxp3^+^, and a vast majority of these expressed CD39. Among the Tconv clusters (Foxp3^−^), two expressed CD39, one of which showed high expression of hallmarks of exhaustion, such as TIM-3, TIGIT, BTLA, CTLA-4, and PD-1 (Tconv CD39+iRs+), while the other barely expressed CTLA-4 or TIM-3 and was negative for TIGIT, BTLA and PD-1 expression (Tconv CD39^+^iRs^−^). Considering that Tconv expressing high levels of iRs and CD39 have been associated with ongoing activation due to chronic antigen exposure at the tumor site ^8^, the heterogeneity within the CD39^+^ T conv may reflect different states of T cell activation.

To characterize the effector ability of human CD39^+^ Tconv cells from tumors and M-dLNs, we evaluated the frequency of TNF, IFNγ and IL-2-producing Tconv upon stimulation. We compared the frequency of cytokine-producing cells between CD39^−^ and CD39^+^ Tconv cells, focusing on non-naïve cells (gating out CD45RA^+^CD27^+^ cells). CD39^+^ Tconv cells from tumors exhibited reduced frequencies of TNF but equal frequencies of IL-2- and IFNγ producing cells, compared to CD39^−^ Tconv cells (Fig. 5E). Also, CD39^+^ Tconv cells from M-dLNs showed conserved production of effector cytokines (Supplementary Fig. S6B). Notably, CD39^+^ Tconv cells from tumors and M-dLNs exhibited higher frequencies of CD107a^+^ cells than their CD39^−^ counterparts (Fig. 5E and Supplementary Fig. S6B).

Altogether, these results indicate that CD39^+^ Tconv cells present in tumors and M-dLNs from BC patients constitute a heterogeneous cell population enriched in activated cells which, although presenting some exhaustion features, are functional in M-dLNs and conserve their cytotoxic potential even under the influence of the TME. Thus, CD39^+^ Tconv cells may emerge as key players in anti-tumor immunity.

### scRNAseq data reveals that CD4-CXCL13 clusters express CD39, iRs and cytolytic markers

To expand our results to other cancers beyond BC, we analyzed available single cell RNA-Seq data of CD4^+^ T cells isolated from tumors of patients with hepatocellular carcinoma (HCC) ^32^, colorectal cancer (CRC) ^33^, and non-small cell lung cancer (NSCLC) ^34^. We found that in CRC, CD39 was expressed in two different clusters of CD4^+^ cells (CD4_C08-IL23R and CD4_C09-CXCL13) besides the three Treg clusters (CD4_C010-FOXP3, CD4_C011-IL10, CD4_C012-CTLA4) (Figure 6A). Notably, the CD4-CXCL13 cluster concomitantly exhibited high expression of exhaustion markers (PD-1, TIGIT, TIM-3, TOX, CTLA4, and BTL-A) and cytotoxic molecules (GZMA, PRF1, and GZMB) (Figure 6B and Supplementary Fig. S7). Furthermore, co-expression of CD39 and iRs was also found in the CD4-CXCL13 cluster in HCC (Figure 6 C, D, and Supplementary Fig. S7). In contrast, CD39 was expressed only in the Treg cluster of non-small cell lung cancer (NSCLC) samples (Figure 6E). Expression of CXCL13 has been previously associated with a tumor neoantigen-specific Tconv subset that can be subdivided into two clusters, one expressing memory and T follicular helper markers, and the other cytolytic markers and iRs ^35^. These results and reported data reinforce the idea that CD39 may be a marker of tumor-specific CD4^+^ T cells.

**Figure 6:**
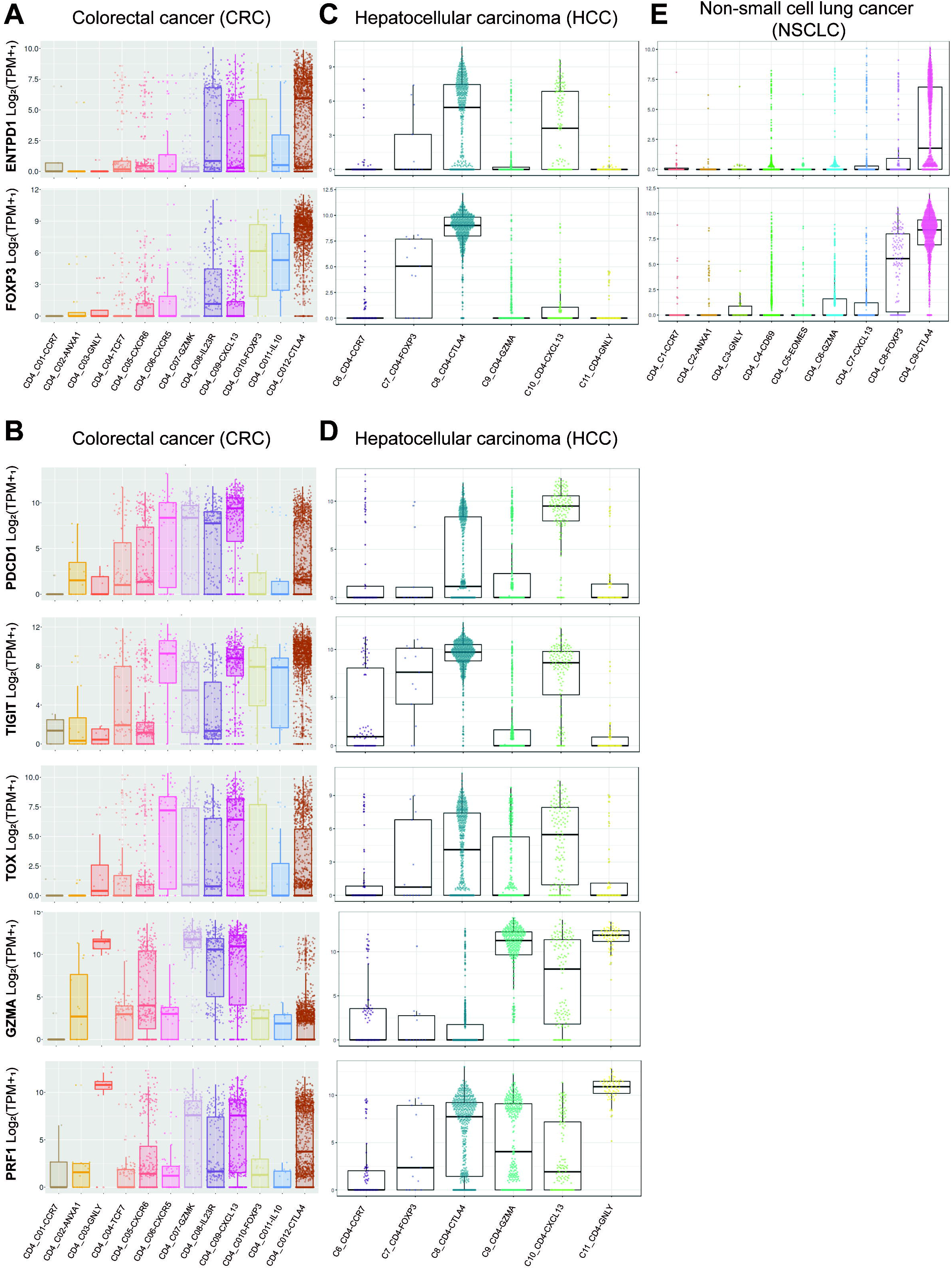
scRNAseq data show that CD39^+^ Tconv cells infiltrate tumors from HCC and CRC patients. **A** Analysis of ENTPD1 (CD39) and FOXP3 expression in single cells isolated from tumors of patients with colorectal cancer (CRC, ^33^). **B** Analysis of PDCD1 (PD-1), TIGIT, TOX, GZMA and PRF1 expression in single cells isolated from tumors of patients with colorectal cancer (CRC, ^33^). **C** Analysis of ENTPD1 (CD39) and FOXP3 expression in single cells isolated from tumors of patients with hepatocellular carcinoma (HCC, ^32^). **D** Analysis of PDCD1 (PD-1), TIGIT, TOX, GZMA and PRF1 expression in single cells isolated from tumors of patients with hepatocellular carcinoma (HCC, ^32^). **E** Analysis of ENTPD1 (CD39) and FOXP3 expression in single cells isolated from tumors of patients with non-small cell lung cancer (NSCLC, ^34^). Clusters are defined in the respective publications. Plots were generated through web-based platforms: http://hcc.cancer-pku.cn/ for HCC; http://lung.cancer-pku.cn/ for NSCLC; http://crctcell.cancer-pku.cn/ for CRC.

### High expression of CD4 and CD39 correlates with increased survival of breast cancer and melanoma patients

To explore the possible contribution of the CD39^+^ Tconv cell subset in the clinical prognosis of BC, melanoma (SKCM), hepatic cancer (LIHC), colon cancer (COAD), rectal cancer (READ), and lung cancer (LUAD), we studied overall survival in a cohort of patients from the TCGA consortium. Samples with high PTPRC (CD45) expression, high CD4 expression, and low FOXP3 expression were selected for analysis. Overall survival was calculated in the selected cohorts, stratifying patients according high or low ENTPD1 (CD39) expression. High gene expression of CD4 and ENTPD1 (CD39) correlated with a better survival of BC (Fig. 7A) and melanoma (Fig. 7B) patients.

**Figure 7:**
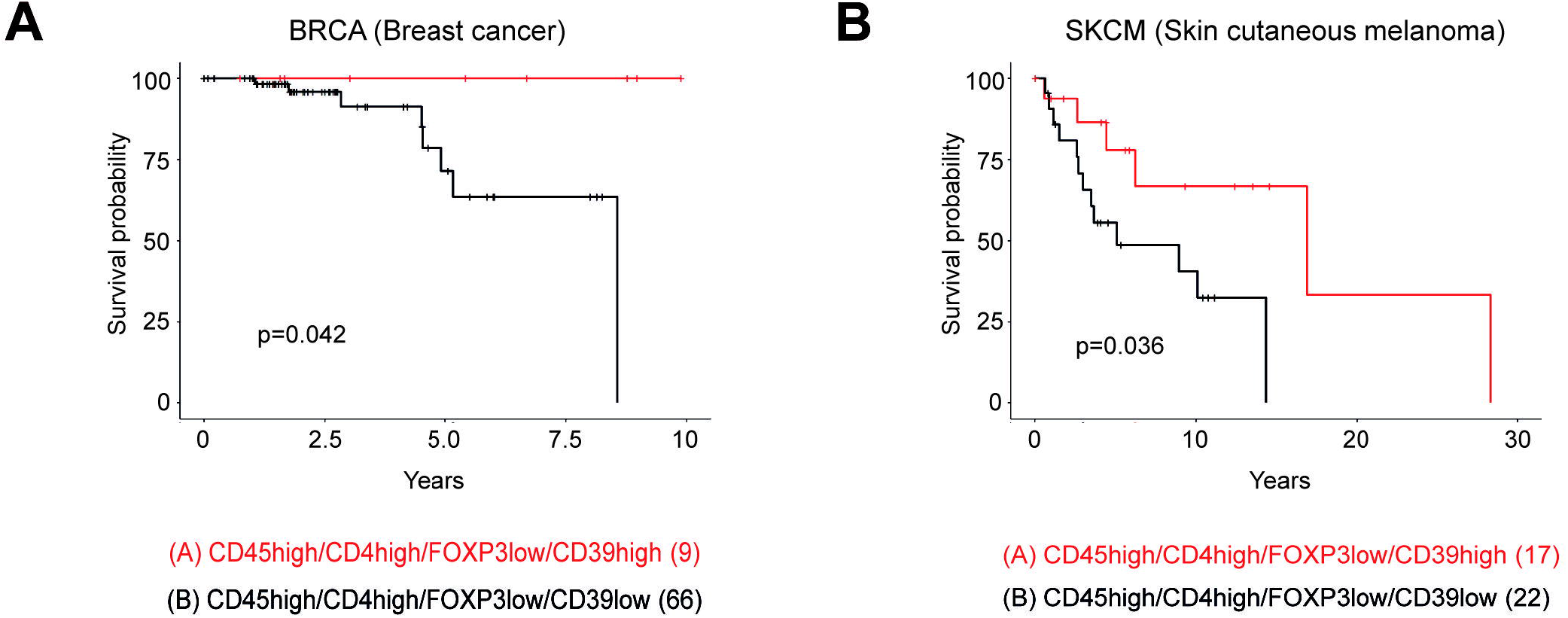
High expression of CD39 correlates with longer survival in patients with BC and melanoma. The overall survival (OS) of patients was calculated according to the expression level of the CD4, PTPRC (CD45), FOXP3 and ENTPD1 (CD39) genes. **A**, TCGA cohort of patients with BC. **B**; TCGA cohort of patients with melanoma. p-values of the survival analyzes were calculated with the log-rank test and are indicated in the graphs. The number of patients in each selected population is indicated in parentheses in the graph.

However, there were no significant differences in the survival of patients with LIHC, COAD, READ and LUAD (Supplementary Fig. S8). This suggests that, at least in some types of cancer, CD39^+^ Tconv are associated with controlling tumor progression and may constitute a key target for tumor immunotherapy.

## Discussion

Cancer immunotherapy has historically focused on CD8^+^ T cells; however, antitumor CD4^+^ T cell immunity has emerged as an attractive new target in recent years ^9^.

Our work demonstrates that a significant frequency of T-I Tconv cells express CD39 in different experimental models. Previously, CD39 has been mostly associated with Treg and used as a marker of this cell subset. Here, we show that another subset of CD4^+^ T cells expressing CD39 comprises a cell population with exhaustion and cytotoxic features. CD39^+^ Tconv cells are present in tumors and M-dLN from BC patients, but appear at lower frequencies or are even absent in PB and NM-dLNs. Their frequency is also higher in tumor than in juxtatumoral breast tissue. These observations emphasize the idea that, in mice and humans, the TME dictates CD39 expression. These results agree with those showing that CD39 expression was significantly elevated in T-I Tconv with respect to circulating CD4^+^ T cells from head and neck squamous cell carcinoma (HNSCC), lung, and colorectal cancer patients ^8, 36^. Similarly, it has been demonstrated that the fraction of CD4^+^ T cells expressing CD39 was also significantly higher in tumor than in healthy lung tissue from NSCLC patients ^37^.

The transcriptional profile as well as multiparametric flow cytometry analysis demonstrated that T-I CD39^+^ Tconv, like its CD39^−^ counterpart, from tumor-bearing mice exhibited an effector memory phenotype, but CD39^+^ Tconv cells expressed multiple iRs. These same iRs have been shown upregulated also in Tconv cells during chronic infections and cancer, and are expressed by exhausted CD8^+^ T cells, suggesting similarities between CD4^+^ and CD8^+^ T cell-exhaustion phenotypes ^19, 30^. Additionally, we found that its cytotoxic potential and capacity to produce IFNγ are other important features of CD39^+^ Tconv. These findings match observations that not all effector functions of CD4^+^ exhausted T cells are necessarily compromised during chronic infection. Thus, perforin and granzyme B production in CD4^+^ T cells was higher in patients infected with HIV than in healthy controls ^38^. Interestingly, such maintenance of cytotoxic functions has also been described for terminally exhausted CD8^+^ T cells ^39^.

Heterogeneity is a hallmark of T cell exhaustion ^15^. For CD8^+^ T cells, the more undifferentiated cells that give rise to the more terminal exhausted ones have been identified as “progenitor exhausted CD8^+^ T cells”, and their transcriptional and regulatory program has been well defined ^40^. However, the TFs guiding exhaustion of T-I CD4^+^ T cells are not yet well identified ^30^. Indeed, different subsets within the exhausted CD4^+^ T cell population remain ill defined. Here, we observed that, compared to CD39^−^ Tconv, CD39-expressing Tconv cells display higher expression of TOX, T-bet, Eomes, Blimp-1 and cMaf, all associated with the acquisition and maintenance of effector functions and progression towards exhaustion ^15^. In addition, concerning TCF-1, a TF associated with a naïve and/or stem-like phenotype, we detected the downregulation of the *Tcf7* transcript as well as lower expression of TCF-1 in CD39^+^ Tconv compared to its CD39^−^ counterpart. Recent studies indicate that T-bet drives the conversion of TCF-1^+^ progenitor exhausted CD8^+^ T cells to TCF-1^−^ cells, promoting the re-engagement of effector functions ^40^. Accordingly, we speculate that high expression of T-bet on CD39^+^ Tconv may be responsible for IFNγ production. Together our results suggest that the exhaustion program of CD4^+^ T cells resembles that of CD8^+^ T cells.

CD39 expression has been associated with antigen-mediated TCR triggering in exhausted CD8^+^ and CD4^+^ T cells. Simoni et al. ^41^ have shown that tumor-specific CD8^+^ T cells from colorectal and lung tumors express CD39. They demonstrated that the lack of CD39 expression could be a marker to identify bystander T-I lymphocytes. Similar observations were made for tumor-specific and bystander CD4^+^ T cells ^8^. In the same direction, Balanca et al. ^19^ investigated antigen specificity in T-I Tconv from ovarian cancer patients, according to PD-1 and CD39 expression. While PD-1^high^CD39^+^ T cells were able to secrete IFNγ upon NY-ESO-1 antigen stimulation, PD-1^high^CD39^−^ did not respond, indicating that T-I PD-1^high^CD39^+^CD4^+^ encompassed tumor antigen-specific cells. More recently, Zheng et al. ^32^ demonstrated that neoantigen-reactive CD4^+^ T cells from tumors of gastrointestinal cancer patients expressed high levels of exhaustion and cytotoxic genes. Based on these results, we hypothesize that T-I CD39^+^ Tconv from BC patients could be tumor-specific.

Currently, cytotoxic CD8^+^ T cells are the focus of efforts to understand how immunotherapy elicits anti-tumor immune responses; however, the importance of heterogeneous subsets of T-I CD4^+^ remains less clear. In our study, we observed that, upon anti-CTLA-4 treatment, MC38 tumor-bearing mice exhibit a considerable expansion of CD39-expressing Tconv in addition to the expected depletion in the Treg population. After anti-CTLA-4 treatment, the expanded CD39^+^ Tconv maintain the expression of molecules and TFs associated with both exhaustion and cytotoxicity, and also upregulate T-bet expression. Higher expression of this TF may prompt these cells to further produce IFNγ, contributing to an improved anti-tumor immune response, as previously reported ^42^. In this line, Liakou et al. ^43^ demonstrated that CTLA-4 blockade increased IFNγ producing ICOS^+^CD4^+^ T cells, shifting the effector to Treg cell ratio in human bladder tumors. More recently, Sledzinska et al. ^44^ demonstrated that increased IL-2 availability after anti-CTLA-4 treatment also contributed to shape the CD4^+^ T cell compartment in the TME.

PD-1 blockade also influences T-I CD4^+^ T cells. Hormone receptor positive or triple negative breast cancer (TNBC) patients treated with anti-PD-1 before surgery exhibited an expansion of CD8^+^ and CD4^+^ T cells irrespective of the tumor subtype. The expanded CD4^+^ T cells were characterized by the expression of Th1 and follicular helper markers. The analysis of DEGs revealed that the expanded CD4^+^ T cells exhibit markers of effector function (IFNG), cytotoxicity (GZMB, PRF1, GZMA), and exhaustion (TOX, TIGIT, PDCD1, LAG3) ^45^. Further studies demonstrated that TNBC patients treated with paclitaxel in combination with anti-PD-1 showed an expansion of CXCL13^+^ T cells (CD4^+^ and CD8^+^) that expressed T-cell exhaustion related genes. Of note, the expansion of CXCL13^+^ T cells correlated with successful response to combination therapy ^33^.

Our analysis of tumor tissue sequencing data from TGCA shows that higher expression of CD4 and CD39 transcripts in samples from BC and melanoma patients correlates with better prognosis. This observation goes in hand with the results obtained by Peter Savas et al. ^46^, confirming a significant association between a tissue-resident memory gene signature (i.e. iRs, CD39 and effector function associated genes) and improved survival in TNBC patients. In addition, CD39^+^CD103^+^PD-1^+^CD8^+^ T-I T cells are associated with prolonged survival in patients with ovarian cancer ^47^. In this regard, one limitation of our analysis is that it does not dismiss the contribution of CD39^+^ expression in CD8^+^ T cells. However, our previous observation that BC tumors exhibit greater proportions of TI-CD4^+^ than CD8^+^ T lymphocytes ^21^ supports the relevance of the contribution of the CD39-expressing CD4^+^ T cell in overall survival.

Our results identify CD39 as a biomarker of CD4^+^ T cells with characteristics of both exhaustion and cytotoxic potential. Uncovering the role of CD39-expressing CD4^+^ T cells in the TME should help design new strategies to improve the efficacy of current immunotherapies.

## Abbreviations

BC: breast cancer
PB: peripheral blood
CTLs: cytotoxic T cells
dLNs: draining lymph nodes
iRs: inhibitory receptors
M-dLNs: metastatic draining lymph nodes
NM-dLNs: non-metastatic draining lymph nodes
OS: overall survival
Tconv: T conventional cells
TFs: transcription factors
T-I: tumor infiltrating
TME: tumor microenvironment
Treg: T regulatory cells.

## Supplementary Legends

**Supplementary Figure S1. Gate strategies used for the analysis of samples from tumor-bearing mice and BC patients. A** Gate strategy used in tumors from tumor-bearing mice. **B** Gate strategy used in dLNs and spleen from tumor-bearing mice. **C** Gate strategy used in tumors and juxtatumoral tissues from BC patients. **D** Gate strategy used in PB from BC patients. **E** Gate strategy used in M-dLNs and NM-dLNs from BC patients.

**Supplementary Figure S2. Tconv cells infiltrating MCA-OVA, MC38 and 4T1 tumors exhibit CD39 expression. A** Representative dot plots and graphs show frequency of CD39^+^ cells gated on Tconv in tumors (red), dLNs (green) and spleen (blue) at day 17 p.i. from MCA-OVA tumor-bearing mice. **B** Representative dot plots and graphs show frequency of CD39^+^ cells gated on Tconv in tumors (red), dLNs (green) and spleen (blue) at days 17 p.i. from MC38-tumor-bearing mice. **C** Representative dot plots and graphs show frequency of CD39^+^ cells gated on Tconv in tumors (red), dLNs (green) and spleen (blue) at days 21 p.i. from 4T1-tumor-bearing mice. All results are representative of 3 independent experiments. Data presented as mean ± SEM. ns, \*\*\*\**p*≤ 0.0001.

**Supplementary Figure S3. Transcriptional profiling of tumor-infiltrating CD4**^**+**^ **T cells. A** Volcano plot shows differentially expressed genes (DEGs) between T-I CD39^+^ Tconv and Treg cells. Genes were considered differentially expressed if FC≥ 2 and adjusted p value<0.05 and these are colored in blue. Upregulated genes are shown in red, downregulated genes in green. Genes of interest are highlighted with their gene symbols. **B** Graph shows selected pathways significantly enriched corresponding to genes upregulated in T-I CD39^+^ Tconv compared to Treg using Panther Classification System. Relevant pathways are colored red.

**Supplementary Figure S4. CD39-expressing Tconv cells express iRs associated with exhaustion**. T-I lymphocytes were obtained from tumors from B16F10-OVA tumor-bearing mice at day 17 p.i. and analyzed by flow cytometry. **A** Line graphs show the expression of iRs related to exhaustion (PD-1, TIGIT, TIM-3, LAG-3 and 2B4) on CD39^+^ (red dots) or CD39^−^ (gray squares) T-I Tconv cells. **B** Line graphs and histograms show the expression of CTLA-4 and PD-L1 on T-I CD39^+^ (red histograms, red dots) or CD39^−^ (gray histograms, gray squares) T-I Tconv cells. All results are representative of 3 independent experiments. **A-B** Lines indicate that data are paired. Paired T test was used to compare CD39^+^ vs CD39^−^ Tconv cells. \**p*≤ 0.05; \*\**P*≤ 0.01.

**Supplementary Figure S5. Treatment with anti-CTLA-4 impacts tumor weight and volume**. (Left panel) Line graph shows the tumor volume from mice injected with MC38 cell line and treated with anti-CTLA-4 (blue) or IgG control (orange). (Right panel) Bar graph illustrates the average tumor weight of mice injected with MC38 cell line and treated with anti-CTLA-4 (blue) or IgG control (orange). Representative results of two independent experiments (5 mice per group). Unpaired t-test and 2-way ANOVA were used to analyze the results. *p≤ 0.05, **p≤ 0.01.

**Supplementary Figure S6. CD39**^**+**^ **Tconv cells are present in M-dLN. A** Line graphs show frequencies of iRs expressing cells within CD39^+^ (red dots) or CD39^−^ (gray squares) Tconv from M-dLNs of BC patients (N=8, 7 or 2). **B** Line graphs show frequencies of cytokine-producing cells (TNF, IFNγ and IL-2) and CD107a^+^ cells in CD39^+^ (red dots) and CD39^−^ (gray squares) Tconv cells from M-dLNs after PMA/Ionomycin stimulation (*N=5-6*). Lines indicate that data are paired. Paired T test was used to compare CD39^+^ vs CD39^−^ Tconv cells. \**p*≤ 0.05.

**Supplementary Figure S7. scRNAseq data show that T-I CD39**^**+**^ **Tconv cells from HCC and CRC patients express iRs and cytotoxic molecules**. Analysis of GZMB, HAVCR2 (Tim-3), BTLA and CTLA-4 expression in single cells isolated from tumors of patients with colorectal cancer (CRC, ^33^) and hepatocellular carcinoma (HCC, ^32^). Clusters are defined in the respective publications. Plots were generated through web-based platforms: http://hcc.cancer-pku.cn/ for HCC; http://lung.cancer-pku.cn/ for NSCLC; http://crctcell.cancer-pku.cn/ for CRC.

**Supplementary Figure S8: Overall survival in patients with different types of cancer**. The overall survival (OS) of patients was calculated according to the expression level of the CD4, PTPRC (CD45), FOXP3 and ENTPD1 (CD39) genes. **A** TCGA cohort of patients with liver hepatocellular cancer (LIHC). **B** TCGA cohort of patients with colon adenocarcinoma (COAD). **C** TCGA cohort of patients with rectum adenocarcinoma (READ). **D** TCGA cohort of patients with lung adenocarcinoma (LUAD). p-values of the survival analyzes were calculated with the log-rank test and are indicated in the graphs. The number of patients in each selected population is indicated in parentheses in the graph.

## Acknowledgments

We thank the staff of Cytometry Core, Animal and Cell culture of CIBICI-CONICET at the Facultad de Ciencias Químicas, UNC, and the Surgery Department of Hospital Rawson, Argentina. We also thank the staff of the Institut Curie Flow Cytometry facility and the Clinical Immunology Laboratory, Institut Curie, for the collection of human samples.

## Author contributions

S.B. conducted experiments, analyzed and interpreted the data, and participated in manuscript writing. C. A., J. T-B., F.P.C. and C.R, participated in experiments with mice and revised the manuscript. M.C.R, C.S., participated in experiments with human samples. W.R., D.R., E.F. and J.T-B. participated in biostatistics and computational analysis. A.V-S, E.B. and A.del C. provided the biological material from cancer patients. A.G. and E.V.A.R. participated in data discussion, interpretation of results and manuscript revision. C.L.M. and E.P., designed and supervised the study, analyzed and interpreted the data and wrote the manuscript.

## Data availability statement

All data relevant to the study are included in the article or uploaded as supplementary information.

