## Supplementary Figures and Tables for "CD39^+^ conventional CD4^+^ T cells with exhaustion traits and cytotoxic potential infiltrate tumors and expand upon CTLA-4 blockade"

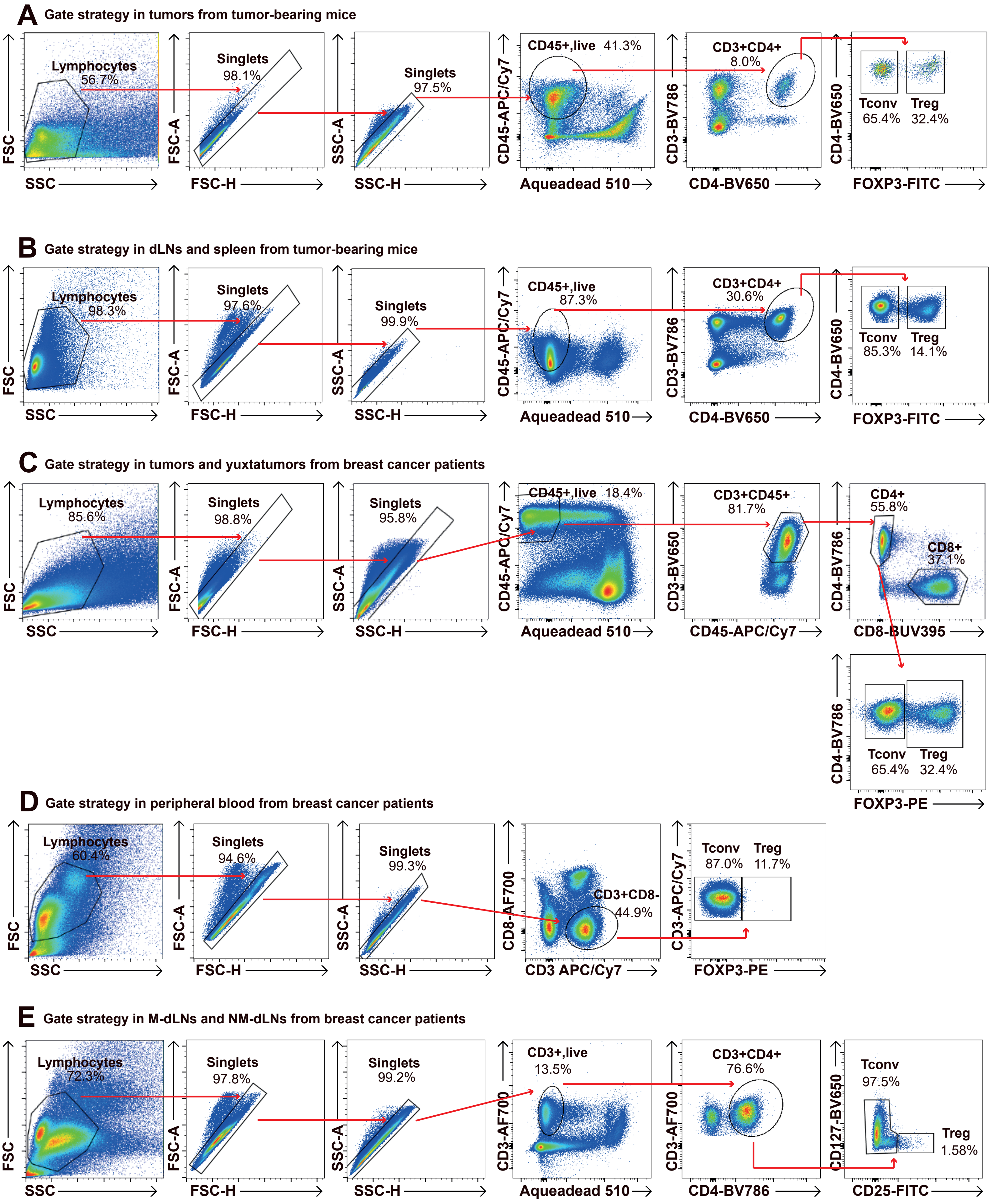
A Gate strategy in tumors from tumor-bearing mice
Lymphocytes 56.7%
Singlets 98.1%
Singlets 97.5%
CD45+,live 41.3%
CD3+CD4+ 8.0%
Tconv 65.4%
Treg 32.4%
FSC
SSC
FSC-A
FSC-H
SSC-A
SSC-H
CD45-APC/Cy7
Aqueadead 510
CD3-BV786
CD4-BV650
FOXP3-FITC
B Gate strategy in dLNs and spleen from tumor-bearing mice
Lymphocytes 98.3%
Singlets 97.6%
Singlets 99.9%
CD45+,live 87.3%
CD3+CD4+ 30.6%
Tconv 85.3%
Treg 14.1%
C Gate strategy in tumors and yuxtatumors from breast cancer patients
Lymphocytes 85.6%
Singlets 98.8%
Singlets 95.8%
CD45+,live 18.4%
CD3+CD45+ 81.7%
CD4+ 55.8%
CD8+ 37.1%
CD3-BV650
CD45-APC/Cy7
CD4-BV786
CD8-BUV395
Tconv 65.4%
Treg 32.4%
FOXP3-PE
D Gate strategy in peripheral blood from breast cancer patients
Lymphocytes 60.4%
Singlets 94.6%
Singlets 99.3%
CD3+CD8- 44.9%
CD8-AF700
CD3 APC/Cy7
Tconv 87.0%
Treg 11.7%
CD3-APC/Cy7
FOXP3-PE
E Gate strategy in M-dLNs and NM-dLNs from breast cancer patients
Lymphocytes 72.3%
Singlets 97.8%
Singlets 99.2%
CD3+,live 13.5%
CD3+CD4+ 76.6%
Tconv 97.5%
Treg 1.58%
CD3-AF700
Aqueadead 510
CD4-BV786
CD127-BV650
CD25-FITC

Figure S1

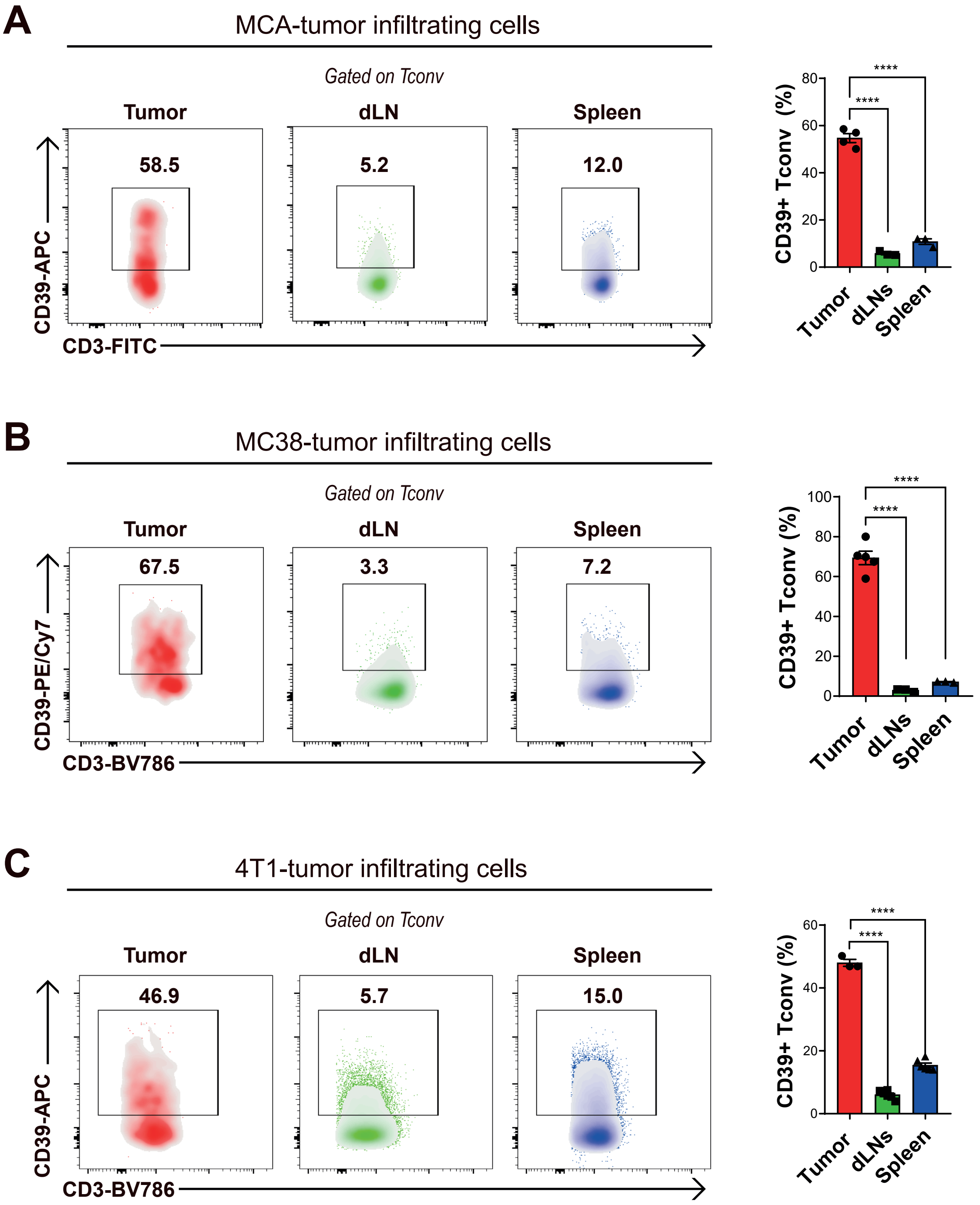
A
MCA-tumor infiltrating cells
Gated on Tconv
Tumor
dLN
Spleen
58.5
5.2
12.0
CD39-APC
CD3-FITC
CD39+ Tconv (%)
****
****
Tumor
dLNs
Spleen
B
MC38-tumor infiltrating cells
Gated on Tconv
Tumor
dLN
Spleen
67.5
3.3
7.2
CD39-PE/Cy7
CD3-BV786
CD39+ Tconv (%)
****
****
Tumor
dLNs
Spleen
C
4T1-tumor infiltrating cells
Gated on Tconv
Tumor
dLN
Spleen
46.9
5.7
15.0
CD39-APC
CD3-BV786
CD39+ Tconv (%)
****
****
Tumor
dLNs
Spleen

Figure S2

A

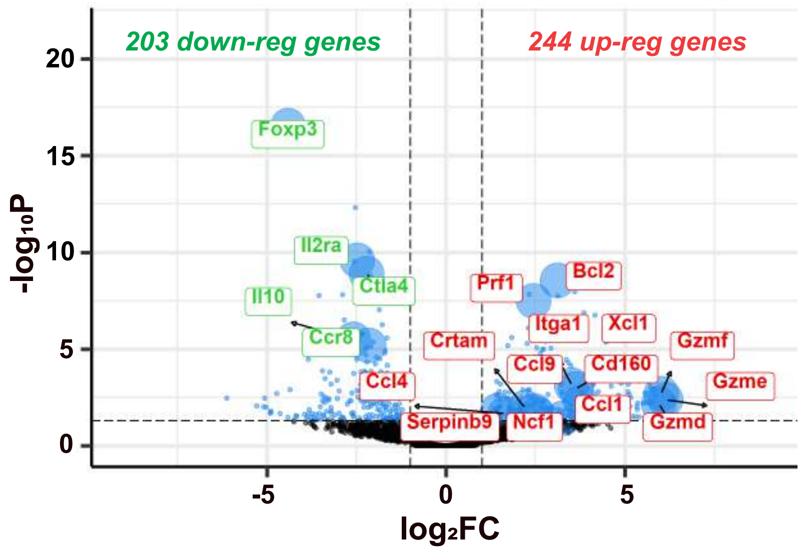

203 down-reg genes
244 up-reg genes
Foxp3
Il2ra
Ctla4
Il10
Ccr8
Ccl4
Serpinb9
Crtam
Prf1
Bcl2
Itga1
Xcl1
Gzmf
Ccl9
Cd160
Gzme
Ccl1
Ncf1
Gzmd
-log10P
log2FC

B

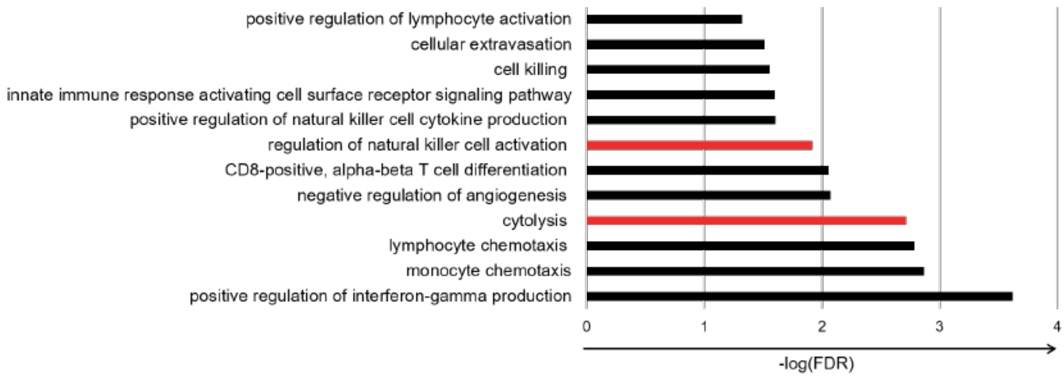

positive regulation of lymphocyte activation
cellular extravasation
cell killing
innate immune response activating cell surface receptor signaling pathway
positive regulation of natural killer cell cytokine production
regulation of natural killer cell activation
CD8-positive, alpha-beta T cell differentiation
negative regulation of angiogenesis
cytolysis
lymphocyte chemotaxis
monocyte chemotaxis
positive regulation of interferon-gamma production
-log(FDR)

Figure S3

A

B

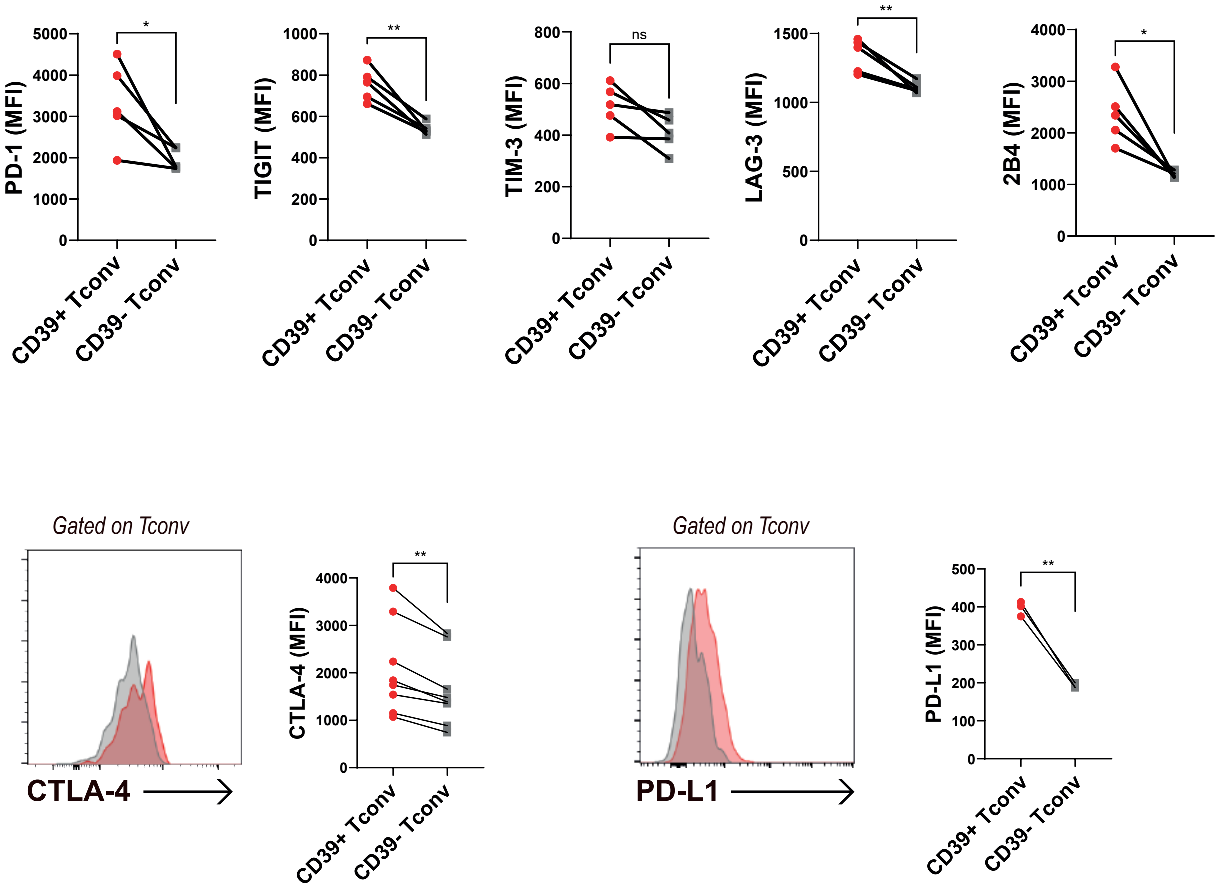

PD-1 (MFI)
TIGIT (MFI)
TIM-3 (MFI)
LAG-3 (MFI)
2B4 (MFI)
CD39+ Tconv
CD39- Tconv
*
**
ns
Gated on Tconv
CTLA-4
CTLA-4 (MFI)
PD-L1
PD-L1 (MFI)

Figure S4

A

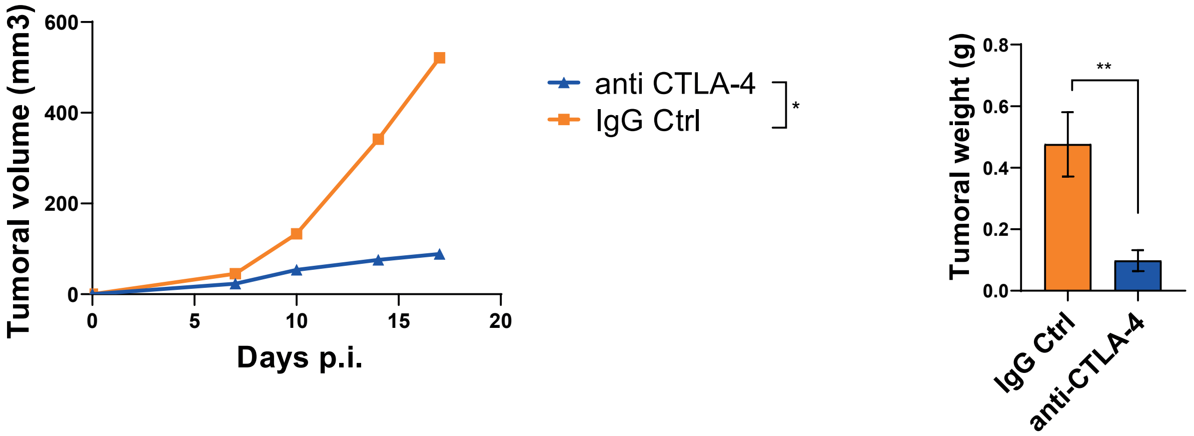

Tumoral volume (mm3)
600
400
200
0
0
5
10
15
20
Days p.i.
anti CTLA-4
IgG Ctrl
*
Tumoral weight (g)
0.8
0.6
0.4
0.2
0.0
**
IgG Ctrl
anti-CTLA-4

Figure S5

**A**

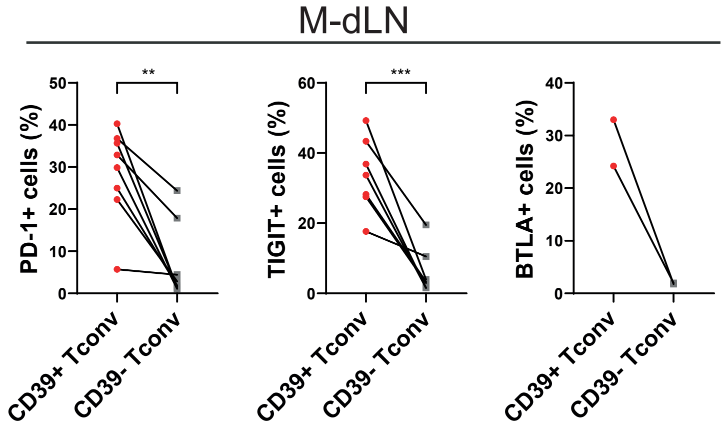
M-dLN
PD-1+ cells (%)
**
TIGIT+ cells (%)
***
BTLA+ cells (%)
CD39+ Tconv
CD39- Tconv

**B**

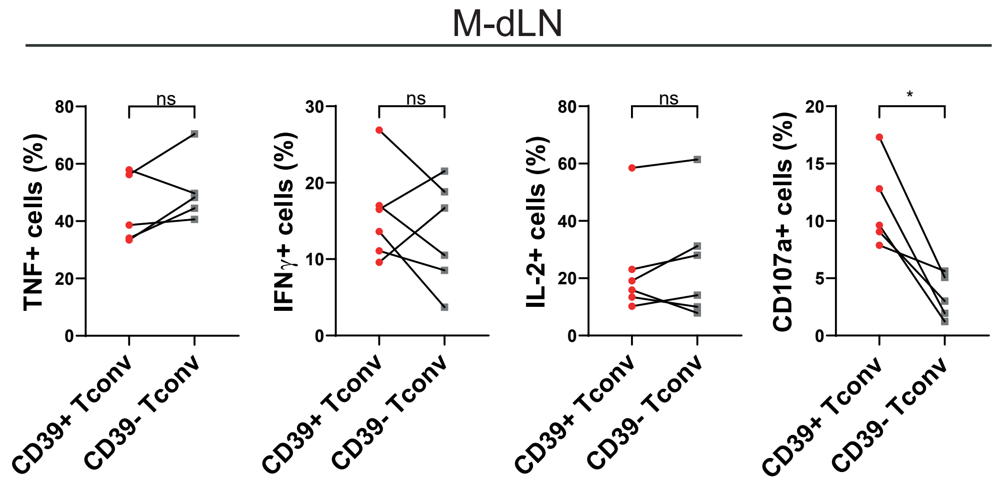
M-dLN
TNF+ cells (%)
ns
IFNγ+ cells (%)
ns
IL-2+ cells (%)
ns
CD107a+ cells (%)
*
CD39+ Tconv
CD39- Tconv

**Figure S6**

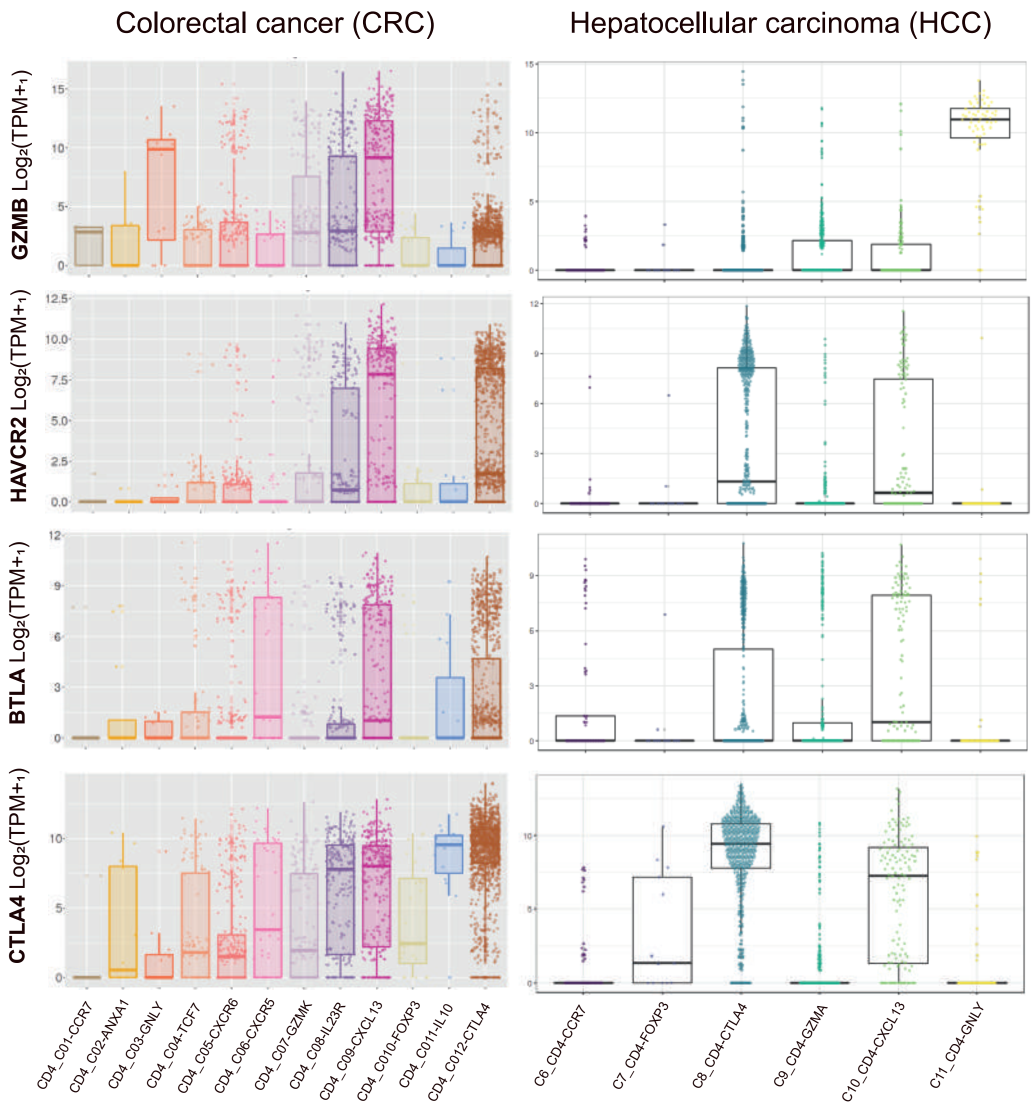

Colorectal cancer (CRC)
Hepatocellular carcinoma (HCC)
GZMB Log2(TPM+1)
HAVCR2 Log2(TPM+1)
BTLA Log2(TPM+1)
CTLA4 Log2(TPM+1)
CD4_C01-CCR7
CD4_C02-ANXA1
CD4_C03-GNLY
CD4_C04-TCF7
CD4_C05-CXCR6
CD4_C06-CXCR5
CD4_C07-GZMK
CD4_C08-IL23R
CD4_C09-CXCL13
CD4_C010-FOXP3
CD4_C011-IL10
CD4_C012-CTLA4
C6_CD4-CCR7
C7_CD4-FOXP3
C8_CD4-CTLA4
C9_CD4-GZMA
C10_CD4-CXCL13
C11_CD4-GNLY

**Figure S7**

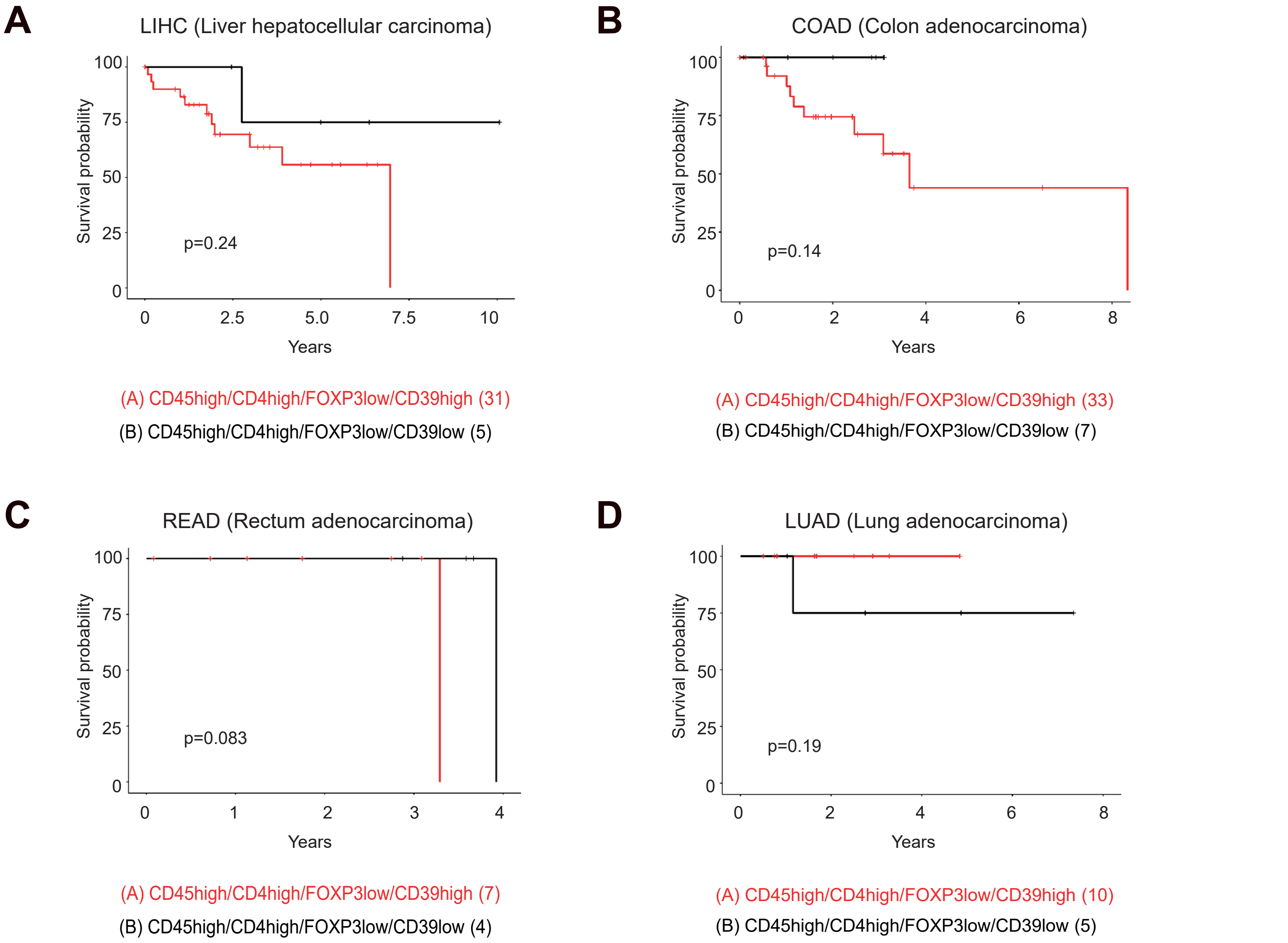
A
LIHC (Liver hepatocellular carcinoma)
Survival probability
100
75
50
25
0
p=0.24
0
2.5
5.0
7.5
10
Years
(A) CD45high/CD4high/FOXP3low/CD39high (31)
(B) CD45high/CD4high/FOXP3low/CD39low (5)
B
COAD (Colon adenocarcinoma)
Survival probability
100
75
50
25
0
p=0.14
0
2
4
6
8
Years
(A) CD45high/CD4high/FOXP3low/CD39high (33)
(B) CD45high/CD4high/FOXP3low/CD39low (7)
C
READ (Rectum adenocarcinoma)
Survival probability
100
75
50
25
0
p=0.083
0
1
2
3
4
Years
(A) CD45high/CD4high/FOXP3low/CD39high (7)
(B) CD45high/CD4high/FOXP3low/CD39low (4)
D
LUAD (Lung adenocarcinoma)
Survival probability
100
75
50
25
0
p=0.19
0
2
4
6
8
Years
(A) CD45high/CD4high/FOXP3low/CD39high (10)
(B) CD45high/CD4high/FOXP3low/CD39low (5)

**Figure S8**

| Clinical and Pathological data of recruited/analyzed Breast Cancer Patients | | |
|---|---|---|
| **Characteristics** | **Patients** | **Value** |
| **Number of patients** | 49 | |
| **Type of sample** | | |
| Tumor | 21 | |
| Juxtatumor | 13 | |
| M-dLN | 10 | |
| NM-dLN | 10 | |
| PB | 5 | |
| **Age (years)** | | 59±13 |
| **Hystology** | | |
| Invasive ductal carcinoma (IDC) | 43 | 88% |
| Invasive lobular carcinoma (ILC) | 4 | 8% |
| Others | 1 | 2% |
| Information not available | 1 | 2% |
| **Stage** | | |
| I | 4 | 8% |
| II | 25 | 51% |
| III | 17 | 35% |
| IV | 0 | 0% |
| Information not available | 3 | 6% |
| **Ki67** | | |
| Positive | 44 | 90% |
| Negative | 0 | 0% |
| Information not available | 5 | 10% |
| **Tumor size (cm)** | | |
| T1 (≤2) | 16 | 33% |
| T2 (2-5) | 27 | 55% |
| T3 (>5) | 5 | 10% |
| Information not available | 1 | 2% |
| **Hormone receptor status** | | |
| Estrogen receptor (ER) + | 39 | 50% |
| Progesterone receptor (PR) + | 35 | 45% |
| HER2 + | 2 | 3% |
| Información no disponible | 2 | 3% |
| **Lymph node status** | | |
| Free (N0) | 18 | 37% |
| N1 (1-3) | 18 | 37% |
| N2 (4-9) | 6 | 12% |
| N3 (>9) | 4 | 8% |
| Información no disponible | 3 | 6% |

Supplementary Table S1

| | Antigen | Clon | Fluorocrome | Brand |
|---|---|---|---|---|
| **Mouse** | 2B4 | eBio244F4 | PE | eBioscience |
| | Blimp-1 | 5E7 | PE-CF594 | BD |
| | CD107a | 1D4B | PE | Biolegend |
| | CD3 | 145-2C11 | APC-Cy7 | BD |
| | CD3 | 145-2C11 | BV785 | Biolegend |
| | CD3 | 145-2C11 | FITC | eBioscience |
| | CD3 | 145-2C11 | PE-Cy7 | eBioscience |
| | CD39 | SolA15 | eFluor 660 | eBioscience |
| | CD39 | 24DMS1 | PE-Cy7 | eBioscience |
| | CD39 | 24DMS1 | PerCP-eFluor 710 | eBioscience |
| | CD4 | GK1.5 | AF700 | Biolegend |
| | CD4 | GK1.5 | Super Bright 645 | eBioscience |
| | CD4 | GK1.5 | PerCP-eFluor 710 | eBioscience |
| | CD4 | RM4-5 | PE-CF594 | BD |
| | CD44 | IM7 | APC-eFluor 780 | eBioscience |
| | CD45 | 30-F11 | AF700 | eBioscience |
| | CD45 | 30-F11 | APC-Cy7 | BD |
| | CD62L | MEL-14 | PE | BD |
| | CD73 | eBioTY/11.8 | PE-Cy7 | eBioscience |
| | CD8 | 53-6.7 | AF700 | eBioscience |
| | C-Maf | Sym0F1 | PE | eBioscience |
| | CTLA-4 | UC10-4B9 | BV605 | Biolegend |
| | Eomes | Dan11mag | PE | eBioscience |
| | Foxp3 | FJK-16s | APC | eBioscience |
| | Foxp3 | RA3-6B2 | FITC | Biolegend |
| | Foxp3 | FJK-16s | PerCP-Cy5.5 | eBioscience |
| | Granzima B | QA16A02 | APC/Fire 750 | Biolegend |
| | Helios | 22F6 | Pacific Blue | eBioscience |
| | IFNg | XMG1.2 | APC | eBioscience |
| | Ki67 | SolA15 | APC | eBioscience |
| | LAG-3 | C9B7W | PerCP-eFluor 710 | eBioscience |
| | PD-1 | 29F.1A12 | BV421 | Biolegend |
| | PD-L1 | MIH5 | PE | eBioscience |
| | Perforina | S16009A | APC | Biolegend |
| | T-bet | eBio4B10 | PE-Cy7 | eBioscience |
| | TCF-1 | S33-966 | PE | BD |
| | TIGIT | GIGD7 | PE-Cy7 | eBioscience |
| | Tim-3 | RMT3-23 | APC | Biolegend |
| | TNF | MP6-XT22 | PE | eBioscience |
| | TOX | TXRX10 | eFluor 660 | Invitrogen |
| **Human** | BTLA | MIH26 | APC | Biolegend |
| | BTLA | J168-540 | PE-CF594 | BD |
| | CCR7 | GO43H7 | BV421 | Ozyme |
| | CD107a | eBioH4A3 | FITC | eBioscience |
| | CD127 | MB15-18C9 | FITC | Miltenyi |
| | CD25 | M-A251 | PE | BD |
| | CD3 | UCHT1 | APC-Cy7 | Biolegend |
| | CD3 | OKT-3 | BV650 | Ozyme |
| | CD39 | eBioA1 | PE-Cy7 | eBioscience |
| | CD39 | A1 | PerCP-Cy5.5 | Biolegend |
| | CD39 | A1 | PE | Biolegend |
| | CD39 | TU66 | BV711 | BD |
| | CD39 | TU66 | Green A 780-60 | BD |
| | CD4 | OKT4 | BV785 | Ozyme |
| | CD45 | 2D1 | APC-Cy7 | BD |
| | CTLA-4 | BNI3 | BV605 | eBioscience |
| | FOXP3 | 236A/E7 | PE | eBioscience |
| | FOXP3 | 236A/E7 | AF488 | eBioscience |
| | IFNg | B7 | BV421 | BD |
| | IFNg | B7 | FITC | BD |

| | | | |
|---|---|---|---|
| IL-2 | MQ1-17H12 | PE | BD |
| PD-1 | J105 | APC | eBioscience |
| PD-1 | J105 | PE-Cy7 | eBioscience |
| PD-1 | EH12.2H7 | BV711 | Ozyme |
| TIGIT | MBSA43 | PerCP-eFluor 710 | eBioscience |
| TNF | MAb11 | APC | eBioscience |

Supplementary Table S2

### DEGs (CD39+ Tconv vs CD39- Tconv)

| UP-reg | FC | UP-reg | FC | UP-reg | FC |
|---|---|---|---|---|---|
| Gzmf | 7,9354 | Lgi2 | 5,2504 | Alox5ap | 4,5525 |
| Filip1 | 7,3080 | Rnase6 | 5,2214 | Lmo2 | 4,5372 |
| Havcr2 | 7,0839 | Clgn | 5,2140 | Siva1 | 4,5333 |
| Rassf4 | 6,9368 | Bend4 | 5,1977 | Mycl | 4,5239 |
| Gldc | 6,8138 | Cd180 | 5,1689 | Il1b | 4,4963 |
| Lyz2 | 6,7516 | Cep112 | 5,1544 | Anpep | 4,4948 |
| Cd244 | 6,7362 | Srgap3 | 5,1056 | Ccl4 | 4,4854 |
| Il10 | 6,6549 | Tgfbi | 5,0947 | 2900026A02Rik | 4,4720 |
| Siglech | 6,3706 | Ccl3 | 5,0781 | Gm17092 | 4,4669 |
| Ccl9 | 6,3559 | Rassf6 | 5,0465 | Gm37881 | 4,4628 |
| Plbd1 | 6,3276 | Mt2 | 5,0308 | Itgam | 4,4581 |
| Mrc2 | 6,3138 | Vash1 | 5,0170 | Bcl11a | 4,4538 |
| Rasd2 | 6,1961 | Tbc1d8 | 4,9603 | Ceacam16 | 4,4327 |
| Gzmc | 6,1165 | Ica1 | 4,9510 | Eldr | 4,4264 |
| Pld4 | 6,0231 | Pirb | 4,9321 | Adgre1 | 4,4196 |
| H2-Eb1 | 6,0063 | Gm43464 | 4,9127 | Pon3 | 4,4123 |
| Klra3 | 5,9756 | Faxc | 4,8809 | Sept3 | 4,4109 |
| Synpo2 | 5,9714 | C1qc | 4,8804 | Rtkn | 4,4000 |
| Csf1r | 5,9379 | Nrgn | 4,8715 | BC030867 | 4,3962 |
| Atp2b2 | 5,8931 | Bag2 | 4,8689 | Ifi207 | 4,3672 |
| Gzmd | 5,8782 | Klra9 | 4,8646 | Slc11a1 | 4,3625 |
| Itgax | 5,8732 | Ifi205 | 4,8584 | Lacc1 | 4,3607 |
| Exoc3l | 5,8128 | Epdr1 | 4,8159 | Ttk | 4,3372 |
| Ltf | 5,8082 | Adamts14 | 4,8052 | Gm5466 | 4,3262 |
| Cacna1e | 5,6593 | Rad54l | 4,7990 | Cd80 | 4,3192 |
| Clec10a | 5,6506 | Aoah | 4,7958 | Tm4sf19 | 4,3098 |
| Clec7a | 5,5916 | Kndc1 | 4,7953 | Gm11223 | 4,2985 |
| Insrr | 5,5731 | Adrb1 | 4,7595 | Sccpdh | 4,2694 |
| Gzme | 5,5410 | Rgs8 | 4,7546 | Etv5 | 4,2630 |
| Lifr | 5,4986 | Mpeg1 | 4,7374 | Napsa | 4,2620 |
| Dscam | 5,4696 | Arnt2 | 4,7254 | Klrc2 | 4,2529 |
| Hist1h2ah | 5,4130 | Tmprss6 | 4,7092 | H2-DMb2 | 4,2525 |
| Blnk | 5,3748 | Cd209a | 4,6876 | Gm14377 | 4,2435 |
| Prf1 | 5,3736 | Gm44174 | 4,6666 | Csf2rb2 | 4,2403 |
| Galnt18 | 5,3343 | Tmem67 | 4,6636 | 3300005D01Rik | 4,2214 |
| Gpld1 | 5,3255 | Rnase4 | 4,6332 | Chl1 | 4,2163 |
| D430041D05Rik | 5,3245 | Chd3os | 4,5771 | Pdgfb | 4,1961 |
| Pkdcc | 5,3233 | Mastl | 4,5706 | Slc7a7 | 4,1445 |
| Fcer1g | 5,3025 | Olfm1 | 4,5562 | Pvrig | 4,1442 |
| Slc17a6 | 5,2761 | Mir24-1 | 4,5545 | D630045J12Rik | 4,1387 |

| UP-reg | FC | UP-reg | FC | UP-reg | FC |
|---|---|---|---|---|---|
| Mcee | 4,1381 | Plac8 | 3,7442 | Tgm2 | 3,1527 |
| Cldnd2 | 4,1311 | Wnt10b | 3,7312 | Cd93 | 3,1517 |
| Setbp1 | 4,1311 | Exph5 | 3,7237 | Spc25 | 3,1432 |
| Gm28513 | 4,1305 | Tmcc3 | 3,7216 | Nusap1 | 3,1403 |
| Gipc3 | 4,1281 | Gins1 | 3,6846 | Cdca2 | 3,1127 |
| Zcchc24 | 4,1243 | Plek | 3,6359 | Trip10 | 3,1017 |
| Dgkg | 4,1177 | Frmd4a | 3,6223 | Nek2 | 3,0903 |
| H2-Ab1 | 4,1116 | Myh10 | 3,6154 | Csf2rb | 3,0783 |
| Slc6a12 | 4,1087 | Wdfy4 | 3,6052 | Birc5 | 3,0665 |
| Wdr95 | 4,1049 | Cdk14 | 3,5996 | Rad51ap1 | 3,0639 |
| Npnt | 4,0964 | Ccr5 | 3,5916 | Bub1 | 3,0567 |
| Hsf4 | 4,0639 | Sirpa | 3,5816 | Spdl1 | 3,0362 |
| Sulf2 | 4,0578 | Itih5 | 3,5426 | Cd160 | 3,0286 |
| Cebpd | 4,0399 | Bspry | 3,5384 | BC055324 | 3,0172 |
| Nfam1 | 4,0235 | Selenon | 3,5380 | Tm6sf1 | 3,0137 |
| H2-Aa | 4,0162 | Hist1h2bh | 3,5363 | Mcm10 | 2,9986 |
| 1700030C10Rik | 4,0119 | Spp1 | 3,5212 | Srxn1 | 2,9863 |
| Ociad2 | 4,0062 | Ccdc184 | 3,5197 | Smpdl3b | 2,9854 |
| Ckap4 | 4,0034 | Asns | 3,5145 | Ccdc34 | 2,9851 |
| Spats2 | 3,9668 | Ildr1 | 3,5102 | Rad51 | 2,9723 |
| Pask | 3,9661 | Plau | 3,4893 | Shcbp1 | 2,9681 |
| Tmem163 | 3,9484 | Cdca5 | 3,4767 | Cdkn2a | 2,9678 |
| Syk | 3,9326 | Mef2c | 3,4761 | Fgr | 2,9363 |
| Stard9 | 3,9224 | Cd74 | 3,4708 | Serpine2 | 2,9315 |
| AA467197 | 3,9138 | Sdf2l1 | 3,4657 | Gcsh | 2,9257 |
| Hist1h4n | 3,8914 | Sapcd2 | 3,4594 | Sgol2a | 2,9146 |
| Ogfrl1 | 3,8698 | Ska1 | 3,4262 | Kdelc2 | 2,9128 |
| Tcrg-C2 | 3,8556 | Esco2 | 3,4145 | Hist1h4j | 2,8902 |
| Cks1b | 3,8331 | Mrps33 | 3,4071 | Pik3ap1 | 2,8664 |
| Galnt3 | 3,8238 | Tnip3 | 3,4044 | Spry2 | 2,8369 |
| Cd8a | 3,8223 | Arhgap33 | 3,3283 | Hist1h2ae | 2,8200 |
| F2rl2 | 3,8218 | Fam71b | 3,3098 | Suox | 2,7790 |
| Sorbs3 | 3,7996 | Tmem256 | 3,3056 | Tnfrsf9 | 2,7657 |
| D630039A03Rik | 3,7967 | Snx9 | 3,2794 | Cep55 | 2,7657 |
| Sgol1 | 3,7924 | Neb | 3,2737 | Adam8 | 2,7648 |
| Zbed3 | 3,7842 | Adssl1 | 3,2448 | Stk38l | 2,7614 |
| Gzmb | 3,7809 | Afdn | 3,2334 | Kif2c | 2,7371 |
| Ccl1 | 3,7629 | Ctnnd2 | 3,2199 | Pclaf | 2,7350 |
| Ciita | 3,7544 | Irf8 | 3,2021 | Adgrg1 | 2,7329 |
| Tlr9 | 3,7519 | Cd8b1 | 3,1970 | Rgs16 | 2,7232 |
| Klrc1 | 3,7510 | Zc3h12c | 3,1808 | Asf1b | 2,7164 |
| Lmtk3 | 3,7482 | Arap3 | 3,1616 | Entpd1 | 2,7147 |

| UP-reg | FC | UP-reg | FC | UP-reg | FC |
|---|---|---|---|---|---|
| Fndc3b | 2,6946 | Cenph | 2,3378 | Cd86 | 2,0658 |
| Klre1 | 2,6885 | Gm6637 | 2,3143 | Tipin | 2,0566 |
| Gsto1 | 2,6790 | Kif14 | 2,3122 | Hist1h3d | 2,0540 |
| Gpr160 | 2,6731 | Ccna2 | 2,3122 | Ticrr | 2,0433 |
| Hist1h2bj | 2,6600 | Cdca8 | 2,2999 | Ptpn5 | 2,0422 |
| Hist1h2ai | 2,6585 | Knl1 | 2,2905 | Fkbp2 | 2,0403 |
| Litaf | 2,6496 | Hist1h2bb | 2,2814 | Ccnf | 2,0400 |
| Cdc45 | 2,6492 | Tdrd7 | 2,2740 | Dgkh | 2,0398 |
| Kifc1 | 2,6449 | Bub1b | 2,2601 | Nek6 | 2,0394 |
| Itgb8 | 2,6423 | Kif22 | 2,2577 | Hist1h2ab | 2,0336 |
| Pi4k2b | 2,6343 | Aurkb | 2,2570 | Tbl2 | 2,0255 |
| Hmmr | 2,6337 | Hist1h4b | 2,2514 | Dusp16 | 2,0169 |
| Anln | 2,6310 | Hist1h2ag | 2,2501 | Hist1h4f | 2,0078 |
| Kif4 | 2,6226 | Hist1h3a | 2,2169 | Spc24 | 2,0005 |
| Lag3 | 2,6143 | Smim3 | 2,2089 | Hist1h2bk | 1,9856 |
| Unc13b | 2,6023 | Neil3 | 2,2021 | Ncaph | 1,9850 |
| Gas2l3 | 2,5688 | Nuf2 | 2,1903 | Ccrl2 | 1,9822 |
| Ckap2 | 2,5648 | Cenpe | 2,1883 | Clspn | 1,9780 |
| Shmt1 | 2,5586 | Klrk1 | 2,1813 | Stmn1 | 1,9742 |
| Xcl1 | 2,5585 | Chst11 | 2,1721 | Ccnb2 | 1,9651 |
| Slc37a2 | 2,5360 | Fignl1 | 2,1655 | Lat2 | 1,9617 |
| Gas7 | 2,5135 | Eomes | 2,1561 | Prdm1 | 1,9584 |
| Hist1h2bg | 2,5062 | Adap1 | 2,1553 | Hist1h2af | 1,9524 |
| 2610318N02Rik | 2,5053 | Aldoc | 2,1425 | Hist1h2bc | 1,9504 |
| Apoe | 2,4875 | Lyn | 2,1420 | Stk32c | 1,9479 |
| Cdk1 | 2,4849 | E2f8 | 2,1389 | Fabp5 | 1,9350 |
| Ncapg | 2,4653 | Tk1 | 2,1378 | Fancd2 | 1,9341 |
| E2f3 | 2,4645 | Srm | 2,1286 | Haus4 | 1,9277 |
| Gpd2 | 2,4642 | Galk1 | 2,1234 | Ccnb1 | 1,9164 |
| Hsd17b7 | 2,4602 | Cdc20 | 2,1205 | Mt1 | 1,9161 |
| Arsb | 2,4576 | C1qtnf6 | 2,1196 | Hist1h3b | 1,9133 |
| Lrrk1 | 2,4425 | Diaph3 | 2,1152 | Prim2 | 1,9079 |
| Itga1 | 2,4387 | Ier5l | 2,1011 | Hist4h4 | 1,8912 |
| Gk | 2,4158 | Hist1h2bm | 2,0993 | Pglyrp1 | 1,8878 |
| Cenpf | 2,4015 | Pdcd1 | 2,0892 | Xdh | 1,8856 |
| Hist1h3i | 2,3988 | Chsy1 | 2,0854 | Gmnn | 1,8801 |
| Zbtb32 | 2,3934 | Pde4a | 2,0842 | Cit | 1,8783 |
| Tubb6 | 2,3867 | Aspm | 2,0790 | Rcn1 | 1,8720 |
| Pcyt1a | 2,3683 | Cdkn2c | 2,0733 | Dut | 1,8701 |
| Klrd1 | 2,3657 | Nkg7 | 2,0718 | Ifi211 | 1,8692 |
| Ckap2l | 2,3609 | Hist1h3c | 2,0678 | Myo1e | 1,8601 |
| Csf1 | 2,3433 | Cst7 | 2,0665 | Knstrn | 1,8480 |

| UP-reg | FC | UP-reg | FC | UP-reg | FC |
|---|---|---|---|---|---|
| Brca1 | 1,8359 | Serpinb6b | 1,5982 | | |
| Hist1h2bn | 1,8126 | Prc1 | 1,5957 | | |
| Bcat1 | 1,8126 | Hist1h1e | 1,5888 | | |
| Hist1h4k | 1,8084 | Hist1h4h | 1,5879 | | |
| Tcf4 | 1,8038 | Serpina3g | 1,5867 | | |
| Hist1h1b | 1,7992 | Sept11 | 1,5845 | | |
| Mki67 | 1,7986 | Fen1 | 1,5814 | | |
| E2f7 | 1,7840 | Kif11 | 1,5808 | | |
| Spag5 | 1,7831 | Nedd4 | 1,5806 | | |
| Serpina3f | 1,7794 | Ybx3 | 1,5798 | | |
| Unc119 | 1,7758 | Tfdp1 | 1,5737 | | |
| Gm45836 | 1,7713 | Smc2 | 1,5686 | | |
| Ptms | 1,7650 | Hist1h2be | 1,5685 | | |
| Cxcr6 | 1,7610 | Mthfd1 | 1,5666 | | |
| Mybl2 | 1,7586 | Chaf1a | 1,5601 | | |
| Top2a | 1,7577 | Zc3h7b | 1,5423 | | |
| Espl1 | 1,7420 | Hist1h1d | 1,5347 | | |
| Lgals3 | 1,7250 | Casp3 | 1,5310 | | |
| Ifng | 1,7222 | Ubash3b | 1,5185 | | |
| Ulbp1 | 1,7211 | Nr4a2 | 1,5165 | | |
| Atad5 | 1,7196 | Rrm2 | 1,5127 | | |
| Dlgap5 | 1,7146 | Tox | 1,5105 | | |
| Hivep3 | 1,7065 | Hist1h4d | 1,4962 | | |
| Ncapg2 | 1,7033 | Serpinb9 | 1,4932 | | |
| Plxnd1 | 1,6998 | Tigit | 1,4658 | | |
| Padi2 | 1,6987 | Coro2a | 1,4525 | | |
| Tpx2 | 1,6978 | Cdk6 | 1,4329 | | |
| Hist1h2an | 1,6968 | Mcm5 | 1,4133 | | |
| Anxa2 | 1,6930 | Hells | 1,4085 | | |
| Kpna2 | 1,6903 | Ncapd2 | 1,3974 | | |
| Hist1h2ac | 1,6839 | Il10ra | 1,3788 | | |
| Kif15 | 1,6816 | Slc2a3 | 1,3463 | | |
| Foxm1 | 1,6784 | Cad | 1,3304 | | |
| Mtbp | 1,6769 | Ccl5 | 1,3261 | | |
| Tuba1c | 1,6755 | Mcm3 | 1,3056 | | |
| Slc16a3 | 1,6717 | | | | |
| Atp8b4 | 1,6704 | | | | |
| Fgl2 | 1,6659 | | | | |
| Stil | 1,6611 | | | | |
| Ehd1 | 1,6441 | | | | |
| Uhrf1 | 1,6358 | | | | |
| Hist1h3e | 1,6093 | | | | |

| DOWN-reg | FC | DOWN-reg | FC | DOWN-reg | FC |
|---|---|---|---|---|---|
| Ssh2 | -1,2191 | Mast4 | -1,7351 | Vipr1 | -2,4420 |
| Cmah | -1,3088 | Klf3 | -1,7654 | Tdrp | -2,4484 |
| Slc12a7 | -1,3356 | Dirc2 | -1,7748 | Gm36931 | -2,5139 |
| Klf2 | -1,3531 | Plcb4 | -1,7864 | Ston1 | -2,5796 |
| Sidt1 | -1,3606 | Gm26740 | -1,7865 | Aff3 | -2,5798 |
| Fam134b | -1,3742 | Nav2 | -1,7947 | E030030I06Rik | -2,5918 |
| Zbtb20 | -1,3962 | Zfp773 | -1,8110 | Ntrk3 | -2,6853 |
| Rasgrp2 | -1,4026 | Lef1 | -1,8373 | Gbp11 | -2,6919 |
| Dgka | -1,4046 | Scml4 | -1,8473 | 5830444F18Rik | -2,7095 |
| Selenop | -1,4118 | Gm11346 | -1,8486 | St8sia6 | -2,7731 |
| Emb | -1,4362 | Gpr146 | -1,8724 | Rasgrf2 | -2,8203 |
| Sesn3 | -1,4387 | Igf1r | -1,8742 | Tnfrsf22 | -2,8224 |
| Fam65b | -1,4397 | Itga7 | -1,8760 | Nsg2 | -2,8583 |
| St8sia1 | -1,4434 | Sestd1 | -1,8924 | Trib3 | -2,8801 |
| Usp28 | -1,4480 | Sh3bp5 | -1,9112 | Gm15675 | -2,8991 |
| Pgap1 | -1,4577 | Treml2 | -1,9408 | Arhgap29 | -2,9421 |
| Kif1b | -1,4795 | Sell | -1,9465 | Cacna1b | -2,9449 |
| Ldlrad4 | -1,4803 | Rapgef4 | -1,9487 | Hdc | -2,9477 |
| Cd55 | -1,4859 | Cbr1 | -1,9634 | Gm12275 | -2,9871 |
| Tnfsf11 | -1,4923 | Thra | -1,9888 | Sfxn5 | -2,9914 |
| Btla | -1,4965 | Tnfsf8 | -1,9960 | Scin | -3,0011 |
| Actn1 | -1,5113 | Gm20186 | -2,0058 | Oaf | -3,0073 |
| Bach2 | -1,5445 | Il9r | -2,0205 | Gm37248 | -3,1158 |
| Il7r | -1,5504 | Map7 | -2,0386 | Acvrl1 | -3,1285 |
| Dtx1 | -1,5545 | Tnfrsf26 | -2,0495 | Gm11168 | -3,1445 |
| Abcg1 | -1,5568 | Il6ra | -2,0819 | Ccr6 | -3,1679 |
| St6gal1 | -1,5617 | Lrig1 | -2,0898 | Cd46 | -3,2213 |
| Bicdl1 | -1,5781 | Utp14b | -2,0976 | Gm11973 | -3,2503 |
| Egr2 | -1,5837 | Stx1a | -2,0994 | G630030J09Rik | -3,4064 |
| Axin2 | -1,5852 | Lmo4 | -2,1313 | Gm17509 | -3,4338 |
| Asap1 | -1,5896 | Tnfrsf25 | -2,1400 | Bambi-ps1 | -3,4343 |
| Nebl | -1,6054 | Tcf7 | -2,1879 | Amigo2 | -3,4357 |
| Patj | -1,6418 | Zbtb11os1 | -2,1978 | Gm44931 | -3,4450 |
| Satb1 | -1,6507 | Rorc | -2,2012 | Mapk12 | -3,4916 |
| Pik3ip1 | -1,6593 | Ccr7 | -2,2066 | Prox2 | -3,5015 |
| Trib2 | -1,6633 | 1810041H14Rik | -2,2920 | Gm45802 | -3,5442 |
| Ttc28 | -1,6700 | Xkr6 | -2,3271 | Gm37802 | -3,5884 |
| Gm26551 | -1,6939 | Dusp8 | -2,3328 | Gm38215 | -3,6099 |
| Trat1 | -1,6987 | Gpr18 | -2,3575 | Hmcn1 | -3,6375 |
| S1pr1 | -1,7146 | Rflnb | -2,3774 | Slc43a1 | -3,6402 |
| Frat2 | -1,7276 | E430014B02Rik | -2,3934 | BC106179 | -3,6917 |
| Slamf6 | -1,7307 | C230085N15Rik | -2,4285 | A530040E14Rik | -3,7080 |

| DOWN-reg | FC |
|---|---|
| Wfikkn2 | -3,7388 |
| Gm10800 | -3,7481 |
| Lama3 | -3,7595 |
| Scn4a | -3,7847 |
| Efemp2 | -3,8318 |
| Gm9889 | -3,9372 |
| Gm26870 | -3,9402 |
| Igfbp4 | -3,9835 |
| Gm10801 | -3,9997 |
| Fcrl1 | -4,0965 |
| Btnl7-ps | -4,1169 |
| Pdlim4 | -4,1497 |
| Gm45778 | -4,1723 |
| Gm17249 | -4,2380 |
| Hmcn2 | -4,2751 |
| Gm14085 | -4,2970 |
| Tspan9 | -4,3295 |
| Ggt5 | -4,3600 |
| Dnhd1 | -4,4186 |
| Ablim3 | -4,5549 |
| Sorcs2 | -4,5628 |
| Gm7967 | -4,6648 |
| Gm44321 | -4,7177 |
| Nr3c2 | -5,0064 |
| Axdnd1 | -5,1678 |
| Sgtb | -5,3067 |
| Tppp | -5,9784 |

### DEGs (CD39+ Tconv vs Treg)

| UP-reg | FC | UP-reg | FC | UP-reg | FC |
|---|---|---|---|---|---|
| Siglech | 6,3508 | Adam11 | 4,8606 | Lifr | 4,1148 |
| Gzme | 6,1379 | Kcnj8 | 4,8573 | Atp2b2 | 4,1014 |
| Gzmf | 6,0128 | Dapl1 | 4,8367 | Samd3 | 4,0688 |
| Gzmd | 5,9467 | Tcrg-C2 | 4,8303 | Slc16a13 | 4,0651 |
| Cpne7 | 5,8438 | Unc13b | 4,8252 | Fcgr2b | 4,0470 |
| Synpo2 | 5,7914 | Cd8a | 4,7003 | Jaml | 4,0340 |
| Gm44174 | 5,7738 | Zbtb16 | 4,6980 | Gm37053 | 4,0232 |
| Gzmk | 5,7149 | D430041D05Rik | 4,6605 | Trbv12-1 | 3,9937 |
| Epha2 | 5,7066 | Hs3st3b1 | 4,6435 | Nrgn | 3,8568 |
| Zfp683 | 5,6840 | Lancl3 | 4,6327 | 1700019D03Rik | 3,8290 |
| Chl1 | 5,5871 | Tcrg-C4 | 4,6229 | 1700025G04Rik | 3,8081 |
| Tgfbr3 | 5,5577 | Bend4 | 4,6136 | Gas7 | 3,8061 |
| Lrrc75b | 5,5173 | Raver2 | 4,5902 | Ncald | 3,7916 |
| S1pr5 | 5,4026 | Fam65c | 4,5797 | Cd244 | 3,7814 |
| Exph5 | 5,3959 | Lgi2 | 4,5738 | Cd160 | 3,7691 |
| Slc17a6 | 5,3550 | Cd63 | 4,5543 | Zfhx3 | 3,7590 |
| Klra9 | 5,3406 | Dlg2 | 4,4965 | Cd40lg | 3,7551 |
| Tdrp | 5,3305 | Tcf7 | 4,4765 | Pawr | 3,7418 |
| Il21 | 5,3031 | Klrc1 | 4,4716 | Tnfsf4 | 3,7330 |
| Slc24a5 | 5,2128 | Wnt10b | 4,4515 | 5830411N06Rik | 3,7031 |
| Trgv2 | 5,1711 | Galnt18 | 4,4461 | Xcl1 | 3,6295 |
| C230096K16Rik | 5,1466 | Exoc3l | 4,4149 | Ly6c2 | 3,6100 |
| Mrc2 | 5,1253 | Gzmc | 4,3930 | Atp8b4 | 3,6030 |
| Klre1 | 5,0961 | Slc6a12 | 4,3846 | Csf2 | 3,5788 |
| Klrc2 | 5,0874 | Arhgef9 | 4,3786 | Plxdc2 | 3,5650 |
| St8sia1 | 5,0718 | Pkd2 | 4,3639 | Pltp | 3,5395 |
| Trbv14 | 5,0648 | Klra3 | 4,3573 | Rnase6 | 3,5195 |
| Ppargc1a | 5,0335 | Fam78b | 4,3177 | Prr5l | 3,5147 |
| Gm44160 | 5,0031 | Eomes | 4,3057 | Itga1 | 3,5072 |
| Epha3 | 4,9959 | Smyd1 | 4,3049 | Fasl | 3,4966 |
| Clec10a | 4,9740 | Tnfsf14 | 4,2635 | Kcnk5 | 3,4939 |
| Eng | 4,9714 | 4933406I18Rik | 4,2473 | Ehd3 | 3,4935 |
| Mgat3 | 4,9687 | Glis1 | 4,2429 | Cdh1 | 3,4772 |
| Qrfp | 4,9654 | Lypd6b | 4,2312 | Pak6 | 3,4743 |
| Pax3 | 4,9577 | Gm26551 | 4,2235 | Hid1 | 3,4729 |
| Ankrd35 | 4,9573 | Faxc | 4,2031 | Frmd4a | 3,4269 |
| Tm4sf19 | 4,9067 | Pde3b | 4,2014 | Sirpa | 3,4232 |
| St3gal6 | 4,8978 | Klrk1 | 4,1659 | Klrd1 | 3,3663 |
| Wdr95 | 4,8933 | Ret | 4,1455 | Nrarp | 3,3583 |
| Ggt1 | 4,8630 | Adgrg1 | 4,1217 | Igfbp4 | 3,3378 |

| UP-reg | FC | UP-reg | FC | UP-reg | FC |
|---|---|---|---|---|---|
| Dmxl2 | 3,3057 | Gldc | 2,5127 | Ms4a4b | 1,8424 |
| Serpine2 | 3,3008 | Ncf1 | 2,4853 | Mirt1 | 1,8160 |
| Atp1b1 | 3,2762 | Lilr4b | 2,4664 | Zadh2 | 1,8019 |
| Ccl1 | 3,2508 | Rapgef4 | 2,4628 | Sh2b3 | 1,7953 |
| Tnfsf8 | 3,2450 | Prf1 | 2,4418 | Tmem163 | 1,7940 |
| Dse | 3,2117 | Ifi204 | 2,4332 | Mfhas1 | 1,7904 |
| Sumf2 | 3,2088 | P2rx7 | 2,4304 | Rasgrp2 | 1,7782 |
| Cd93 | 3,1426 | Ctnnd2 | 2,4068 | Btbd11 | 1,7687 |
| Klf2 | 3,1265 | Tcf4 | 2,4051 | Slco4a1 | 1,7462 |
| Cacna1e | 3,1263 | Adamts14 | 2,3324 | RP23-353P23.2 | 1,7413 |
| Clgn | 3,1218 | Themis | 2,3319 | Il18rap | 1,7352 |
| Bcl2 | 3,1104 | Pde7a | 2,3067 | Abcb1b | 1,7009 |
| 4930513N10Rik | 3,0708 | Dzip1 | 2,2920 | Enc1 | 1,6984 |
| Nqo2 | 2,9788 | Trbv13-1 | 2,2904 | Ccl4 | 1,6583 |
| Cd8b1 | 2,9333 | Ifng | 2,2872 | Itga4 | 1,6420 |
| Nrp2 | 2,9255 | Hdgfrp3 | 2,2829 | Fosb | 1,5324 |
| Ccl6 | 2,9017 | Pld4 | 2,2810 | Slamf7 | 1,5312 |
| Gm26802 | 2,8912 | Pde11a | 2,2793 | Cd226 | 1,5201 |
| Enpp2 | 2,8638 | Ccl3 | 2,2698 | Prkd3 | 1,5199 |
| Parp8 | 2,8525 | Gm14029 | 2,2380 | Gnptab | 1,5136 |
| Rgs8 | 2,8471 | Rflnb | 2,2378 | Asap1 | 1,5134 |
| Slc25a23 | 2,8130 | Crtam | 2,2079 | Actn1 | 1,5014 |
| Adrb1 | 2,8008 | Prag1 | 2,2052 | Hspa1a | 1,4950 |
| Evl | 2,7837 | Tmem238 | 2,1653 | Egr1 | 1,4930 |
| Gm37387 | 2,7763 | Pou6f1 | 2,1540 | Ctsw | 1,4715 |
| Rtn4rl1 | 2,7671 | Mef2c | 2,1323 | Dennd2d | 1,4567 |
| Lair1 | 2,7581 | Mpeg1 | 2,1176 | Slc20a1 | 1,4526 |
| Spry2 | 2,7457 | Nek6 | 2,0802 | Serpinb6b | 1,4441 |
| Mctp2 | 2,7251 | Tbkbp1 | 2,0773 | Pik3r5 | 1,4430 |
| Nedd4 | 2,7125 | Nebl | 2,0576 | Serpinb9 | 1,4411 |
| Arl4d | 2,6837 | Cd7 | 2,0459 | Ugcg | 1,4323 |
| Dtx4 | 2,6729 | Clec2i | 2,0457 | Hspa1b | 1,4298 |
| Dtx1 | 2,6615 | Rhob | 2,0354 | B4galnt1 | 1,4221 |
| Klf3 | 2,6362 | Itgb1 | 2,0254 | Cd96 | 1,4022 |
| Nod1 | 2,5850 | Chn2 | 2,0114 | Lyst | 1,4016 |
| Vipr1 | 2,5708 | Ust | 2,0065 | Arid5a | 1,3503 |
| Ccl9 | 2,5487 | Dock5 | 1,9553 | Syne2 | 1,3425 |
| Tbc1d4 | 2,5470 | Cxcr6 | 1,9326 | Zfp652 | 1,3409 |
| Gpd2 | 2,5451 | Gramd3 | 1,9268 | Dennd4c | 1,3246 |
| Treml2 | 2,5394 | Aff3 | 1,9228 | Ccdc50 | 1,3146 |
| Sema4a | 2,5247 | AB124611 | 1,8939 | | |
| Bach2 | 2,5145 | Smpdl3b | 1,8823 | | |

| DOWN-reg | FC | DOWN-reg | FC | DOWN-reg | FC |
|---|---|---|---|---|---|
| Lrba | -1,2102 | Tnfrsf18 | -1,6394 | Vaultrc5 | -2,0479 |
| Cd2 | -1,2357 | Gm22513 | -1,6446 | Itga3 | -2,0647 |
| Il12rb1 | -1,2493 | Ttn | -1,6456 | Fam124b | -2,0776 |
| Ptprv | -1,2603 | Slc52a3 | -1,6595 | Ebi3 | -2,0861 |
| Trib1 | -1,2796 | Tmbim1 | -1,6659 | Acot11 | -2,0911 |
| N4bp1 | -1,2905 | Gm45266 | -1,6790 | Kif13a | -2,0926 |
| Dusp4 | -1,2951 | Hemk1 | -1,6833 | Lamc1 | -2,0950 |
| Zfp36l1 | -1,3097 | Eif4e3 | -1,6885 | Gm6637 | -2,0967 |
| Rarg | -1,3406 | Sell | -1,6925 | Samd11 | -2,0971 |
| Trp53i11 | -1,3500 | Capg | -1,6963 | Ramp1 | -2,1162 |
| Ctsz | -1,3546 | Ndrg1 | -1,6983 | Tnfrsf4 | -2,1327 |
| Wls | -1,3551 | Gm2a | -1,7083 | Rapsn | -2,1524 |
| Myo1e | -1,3640 | Cpd | -1,7256 | Gng12 | -2,1563 |
| Tspan32 | -1,3769 | Tjp3 | -1,7261 | St3gal5 | -2,1568 |
| Phlpp1 | -1,3893 | Vav2 | -1,7335 | Ccr8 | -2,1574 |
| Icos | -1,3958 | Coq8a | -1,7344 | Arrdc4 | -2,1670 |
| Tnfrsf1b | -1,4039 | Iigp1 | -1,7381 | Angptl2 | -2,2037 |
| Ighm | -1,4063 | Bnip3 | -1,7426 | Ctla4 | -2,2148 |
| Itgb7 | -1,4088 | Slc15a3 | -1,7650 | Rny1 | -2,2156 |
| Zc3h12d | -1,4124 | Isg20 | -1,7712 | Gm25360 | -2,2251 |
| Tnfrsf9 | -1,4151 | Ccr2 | -1,7802 | Sv2c | -2,2343 |
| Nt5e | -1,4158 | Gatsl2 | -1,7990 | Izumo1r | -2,2344 |
| Tmem154 | -1,4395 | Snord118 | -1,8031 | Gm25099 | -2,2363 |
| Sh3pxd2a | -1,4438 | Ass1 | -1,8141 | Cd81 | -2,2378 |
| Aldoc | -1,4500 | Src | -1,8264 | Arhgef5 | -2,2426 |
| Tigit | -1,4808 | RP24-233B16.8 | -1,8511 | Gm28112 | -2,2667 |
| Penk | -1,4980 | Glrx | -1,8522 | Unc13a | -2,2757 |
| Nfil3 | -1,4993 | Gm8818 | -1,8526 | Katnal1 | -2,2853 |
| Rasgef1b | -1,5267 | Bmyc | -1,8535 | Cd72 | -2,3037 |
| Cysltr2 | -1,5327 | Fgl2 | -1,8650 | Gpr55 | -2,3367 |
| Dst | -1,5338 | Gm25939 | -1,8794 | Ikzf4 | -2,3588 |
| Timp2 | -1,5470 | Ptpn5 | -1,8884 | Rgs9 | -2,3626 |
| Ptgfrn | -1,5528 | Axl | -1,9042 | Matk | -2,3636 |
| H1f0 | -1,5587 | Gm22973 | -1,9069 | F830016B08Rik | -2,3700 |
| Rnu12 | -1,5644 | Fam83g | -1,9129 | Klrg1 | -2,3794 |
| Ets2 | -1,5755 | Tlr1 | -1,9372 | Ctif | -2,3876 |
| Psen2 | -1,5824 | Nol4l | -1,9391 | Areg | -2,4013 |
| Sh3rf1 | -1,5833 | Snx9 | -1,9410 | Oit3 | -2,4067 |
| Cep85l | -1,5911 | Swap70 | -1,9639 | Tbc1d30 | -2,4312 |
| Crlf2 | -1,6117 | Arl5a | -1,9791 | Myo1h | -2,4594 |
| Gm4951 | -1,6161 | Tnfrsf13b | -2,0166 | Il2ra | -2,4780 |
| Maf | -1,6339 | Plxnd1 | -2,0297 | Ankrd6 | -2,4822 |

| DOWN-reg | FC | DOWN-reg | FC | DOWN-reg | FC |
|---|---|---|---|---|---|
| Il1r2 | -2,4942 | Lamc2 | -3,4450 | | |
| Ikzf2 | -2,5247 | Vill | -3,4522 | | |
| Neb | -2,5472 | Prune2 | -3,4784 | | |
| n-R5s193 | -2,5666 | Nos1 | -3,4801 | | |
| Gm37347 | -2,5729 | Matn2 | -3,5115 | | |
| Ptgs1 | -2,5828 | 2210408F21Rik | -3,5294 | | |
| Il10 | -2,5869 | Itgae | -3,5454 | | |
| Ppp1r3fos | -2,5999 | Zan | -3,5677 | | |
| Cep112 | -2,6174 | Ppp4r4 | -3,6096 | | |
| Ccr3 | -2,6356 | Trav13-1 | -3,6234 | | |
| Wisp1 | -2,6497 | Gpr45 | -3,6853 | | |
| Gm29113 | -2,6918 | Pcsk1 | -3,7284 | | |
| Epcam | -2,7525 | Irf6 | -3,7558 | | |
| n-R5s155 | -2,8335 | Gm45206 | -3,8217 | | |
| Lrrc32 | -2,8431 | Ncmap | -3,8353 | | |
| Gm10800 | -2,8792 | Zfp111 | -3,8471 | | |
| Itgb8 | -2,8855 | Gm38157 | -3,8529 | | |
| Piwil2 | -2,9045 | Gm14295 | -3,8883 | | |
| Cdcp1 | -2,9346 | RP23-349H12.3 | -3,9456 | | |
| Fam81a | -2,9515 | Slc30a2 | -4,1665 | | |
| Tnfrsf8 | -2,9857 | Gm42436 | -4,1882 | | |
| Pard6g | -2,9867 | Trav1 | -4,2390 | | |
| Muc4 | -3,0295 | Ankrd55 | -4,3557 | | |
| Gm29112 | -3,0434 | Slc24a3 | -4,4080 | | |
| Gm26870 | -3,0964 | Foxp3 | -4,4164 | | |
| Ephx1 | -3,1185 | Gm44321 | -4,6287 | | |
| Hc | -3,1907 | Msrb3 | -4,7194 | | |
| Gm10801 | -3,2032 | 2010107G23Rik | -4,8917 | | |
| Gm38009 | -3,2087 | Btnl5-ps | -4,8920 | | |
| Igsf23 | -3,2113 | 4933407L21Rik | -4,8978 | | |
| Abcc3 | -3,2239 | Gm13522 | -5,0268 | | |
| Nid2 | -3,2390 | Magi1 | -5,0539 | | |
| Ky | -3,2704 | Dnaaf3 | -5,0977 | | |
| Stab1 | -3,2845 | Tmc3 | -5,2841 | | |
| Gm42731 | -3,3290 | Kcnj15 | -6,1243 | | |
| 9430020K01Rik | -3,3350 | | | | |
| Smim10l2a | -3,3567 | | | | |
| Rln3 | -3,3723 | | | | |
| Col18a1 | -3,3910 | | | | |
| Apol9b | -3,3943 | | | | |
| Muc3a | -3,4129 | | | | |
| Hap1 | -3,4227 | | | | |

Supplementary Table S4
